# Concatemer Assisted Stoichiometry Analysis (CASA): targeted mass spectrometry for protein quantification

**DOI:** 10.1101/2024.07.26.605382

**Authors:** Jiaxi Cai, Yun Quan, Cindy Yuxuan Zhang, Ziyi Wang, Stephen M. Hinshaw, Huilin Zhou, Raymond T. Suhandynata

## Abstract

Large multi-protein machines are central to multiple biological processes. However, stoichiometric determination of protein complex subunits in their native states presents a significant challenge. This study addresses the limitations of current tools in accuracy and precision by introducing concatemer-assisted stoichiometry analysis (CASA). CASA leverages stable isotope-labeled concatemers and liquid chromatography parallel reaction monitoring mass spectrometry (LC-PRM-MS) to achieve robust quantification of proteins with sub-femtomole sensitivity. As a proof-of-concept, CASA was applied to study budding yeast kinetochores. Stoichiometries were determined for *ex vivo* reconstituted kinetochore components, including the canonical H3 nucleosomes, centromeric (Cse4^CENP-A^) nucleosomes, centromere proximal factors (Cbf1 and CBF3 complex), inner kinetochore proteins (Mif2^CENP-C^, Ctf19^CCAN^ complex), and outer kinetochore proteins (KMN network). Absolute quantification by CASA revealed Cse4^CENP-A^ as a cell-cycle controlled limiting factor for kinetochore assembly. These findings demonstrate that CASA is applicable for stoichiometry analysis of multi-protein assemblies.

**Summary:** This study presents Concatemer-Assisted Stoichiometry Analysis (CASA) to address a common challenge in cell biological research: quantifying the number of each protein subunit in a native protein complex.

## Introduction

Stoichiometric analyses and absolute quantifications of native protein complexes have been challenging considering the complexity of native biological samples and the precision required to confidently determine protein ratios within 2-fold (e.g., 3:2 or 4:3). To address this, we present concatemer assisted stoichiometry analysis (CASA) and its application to study one of the most intricate multi-protein assemblies of the cell — the kinetochore.

In eukaryotes, kinetochores ensure the faithful segregation of chromosomes during cell division and act as the load-bearing junctions between centromeric chromatin and spindle microtubules. Despite considerable protein sequence divergence across the eukaryotic kingdoms, kinetochores have a broadly conserved structural organization and dozens of functionally conserved subunits [1,2]. Kinetochores comprise two sub-regions: the inner and outer kinetochores, each containing multiple protein sub-complexes. Inner kinetochore proteins assemble on chromosomal centromeres, while outer kinetochore proteins build upon the inner kinetochore and bind microtubules. Because of this structural and functional conservation, budding yeast *Saccharomyces cerevisiae* has been extensively used as a model organism for characterizing kinetochore structures and functions. Prior studies have identified most, if not all, kinetochore subunits in yeast [3–12]. Structures of many kinetochore sub-complexes have also been determined [13–23]. Moreover, dynamic interactions between centromeres, kinetochore proteins, and microtubules have been examined through fluorescent microscopy [24–29].

Despite these advances, the budding yeast kinetochore has yet to be fully reconstituted using recombinant proteins. For instance, Mif2, the budding yeast ortholog of human CENP-C, is essential for kinetochore assembly [9,10,30] but has yet to be included when reconstituting kinetochores *in vitro* [31]. Post-translational modifications of Mif2^CENP-C^ control inner kinetochore assembly [32–34]; however, our understanding of the regulatory mechanism is still incomplete. Cse4, the yeast ortholog of human CENP-A, is a histone H3 variant specific to centromeric chromatin [1,13]. Despite reports of biochemical Cse4-centromere reconstitutions [18,35], capturing this complex in quantities suitable for structural or mechanistic studies has been a major challenge. Recent reported efforts include the use of a single-chain antibody fragment [23] and/or the use of a chimeric Centromere III (CEN III) and the Widom-601 fusion DNA fragment [31] to stabilize the association between the Cse4^CENP-A^-containing histone octamer, CEN DNA, and the essential CBF3 complex. Perhaps this is not surprising, considering that the chaperone Scm3^HJURP^ is required for assembling the Cse4^CENP-A^-nucleosome in cells, along with the CBF3 complex that directly recognizes the native centromere [36–41]. Therefore, the mechanism and end-state product describing the assembly of the Cse4^CENP-A^-nucleosome on native centromeres have yet to be fully understood.

Considerable progress has been made towards reconstituting kinetochores on native centromeric DNA by leveraging the well-defined DNA sequences of budding yeast’s point centromere in conjunction with concentrated yeast whole-cell extracts [36,42–44]. The *ex vivo* kinetochore reconstitution system mirrors the physiological requirements for native kinetochore assembly, including the conserved CDE III sequence and the presence of the CBF3 complex [42]. Single-molecule studies have begun to reveal quantitative aspects of *ex vivo* reconstituted kinetochores [45]. However, kinetochore subunit stoichiometry in this system has yet to be determined due to the low abundance of kinetochores assembled via this approach. The kinetochore problem highlighted the need for sensitive and robust protein quantification in biological samples, inspiring the development of concatemer assisted stoichiometry analysis (CASA), a targeted LC-MS/MS protein quantification platform.

LC-MS/MS has been widely used to analyze peptides and proteins. In most cases, untargeted LC-MS/MS approaches are used to identify and quantify peptides/proteins, particularly when peptides/proteins of interest are unknown [46]. When the analytes of interest are known, targeted LC-MS/MS approaches such as Multiple-Reaction-Monitoring (MRM) and Parallel-Reaction-Monitoring (PRM) have been shown to provide superior selectivity and sensitivity [47–52]. Improvements in quantitative accuracy and precision are especially apparent when targeted LC-MS/MS approaches are combined with isotope dilution using stable isotope-labeled internal standard [53,54]. There are several approaches for generating stable isotope-labeled internal standard to facilitate absolute quantification of proteins, such as synthesis of isotope-labeled peptides, metabolic protein labeling, and quantitative concatemers (i.e., QconCAT) [55]. Specifically, quantitative concatemers have been successfully adopted for biological applications to quantify multiple proteins in a single experiment [56], making them ideal for monitoring larger protein complexes.

To develop a method useful for quantifying subunit abundances in a single large protein complex, a kinetochore-specific concatemer was produced by stable isotope labeling by amino acids in cell culture (SILAC) [57] and purified from a budding yeast expression system. For simplicity, this concatemer is referred to as the concatenated kinetochore protein (CKP). The CKP contained tryptic peptides derived from over 20 kinetochore subunits and was used as stable isotope-labeled internal standard for a targeted liquid chromatography parallel reaction monitoring mass spectrometry (LC-PRM-MS) method. The analytical performance of CASA for kinetochore was evaluated, revealing its linearity, sensitivity, accuracy, and precision. CASA was then used to study *ex vivo* reconstituted kinetochores as a proof-of-concept. The development and application of CASA not only serve the purpose of validating and expanding our existing knowledge of the yeast kinetochore but also hold the potential for facilitating accurate quantitative analysis of other protein complexes.

## Material and methods

### The concatenated kinetochore protein (CKP) construct

For kinetochore subunits (listed in Table 1), peptide candidates for targeted MS were identified following the analysis by data-dependent acquisition LC-MS/MS of immunoprecipitations of Ame1, Mif2, Cse4, and Ndc80 using Ame1-TAF (HZY2464), Mif2-TAF (HZY2347), 3xFLAG-Cse4 (HZY2777), and Ndc80-TAF (HZY2461) strains (see **S1 Table** and purification results are shown in **S1 Figure**). Trans-Proteomic Pipeline (TPP, Seattle Proteome Center) [58] v6.3.2 Arcus was used to analyze MS data from the Saccharomyces Genome Database (SGD, Stanford University) as described previously [59,60]. Briefly, MS data were searched using COMET, and peptides were quantified using XPRESS in label-free mode. For database searching, a static modification of 57.0215 Da was added for carboxy-amido-methylation of cysteine residues, and a differential modification of 15.9949 Da was added for oxidized methionine residues. The final list of targeted peptides was selected from the kinetochore peptides identified by this search (**S2 Table**) based on the integrated signal intensity of the precursor ion, using the following criteria: 1) between 7 and 15 amino acid residues, 2) no methionine or cysteine residues, 3) being fully tryptic peptide, 4) ending with arginine, and 5) being unique in the yeast proteome. Quality control was performed for peptide identification through manual inspection of chromatography and MS/MS spectral assignments. A gene block was then designed as follows: An N-terminal flanking sequence of 5’ – AATCTATATTTTCAAGGTGGATCCACTAGTTCTAGA – 3’ (homology to the TEV cleavage site sequence in the plasmid HZE3236), followed by the sequences of the tryptic peptides lined up head to tail, and finally by a C-terminal flanking sequence of 5’ – GGGGGTTCTCATCATCATCATCATCATGGGGGCGGA – 3’ (homology to the 6xHis-3xFLAG sequence in HZE3236). The gene block was codon optimized and obtained from Integrated DNA Technologies, USA. HZE3236 plasmid linearized by NotI (Cat #: R3189; NEB, England) was repaired by the gene block via homologous recombination in *S. cerevisiae* to make HZE3361. HZE3361 was rescued from yeast genomic DNA by electroporation using electro-competent *E. coli* cells (in-house made). A single colony from the rescue was picked and grown in 5 mL Lysogeny Broth (LB) with 100 µg/mL ampicillin to saturation, pelleted, and mini-prepped (GeneJET Plasmid Miniprep Kit, Cat #: K0503; Thermo Fisher, USA) to obtain the HZE3361 plasmid. HZE3361 was transformed into the yeast strain SCY249 to create yeast strain HZY3059 (**S1 Table**). Galactose induction from this strain led to the expression of the CKP containing a glutathione-S-transferase (GST from *Schistosoma japonicum*) tag at the N-terminus and a 6xHis-3xFLAG tag at the C-terminus.

**Table 1.**
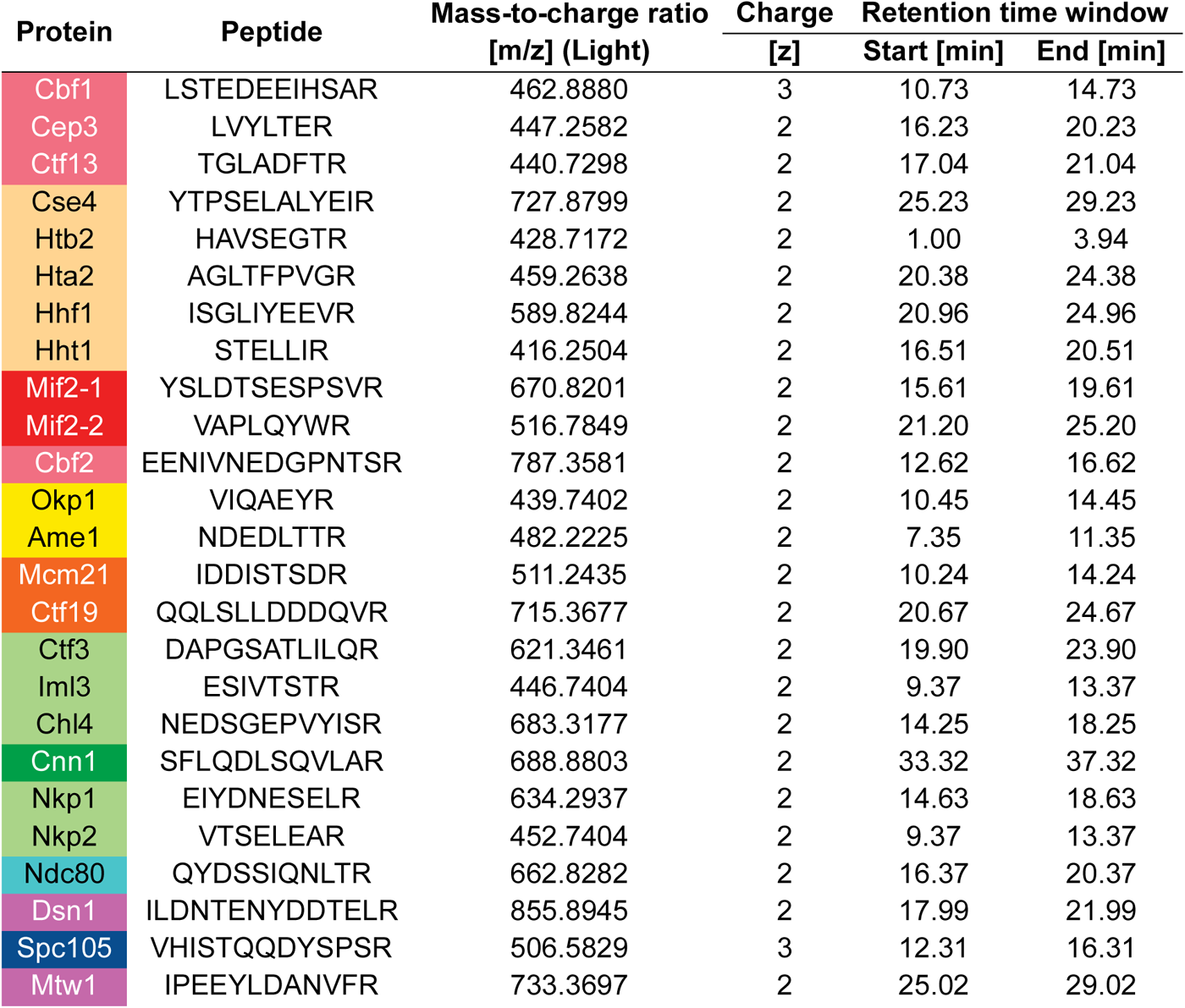
Peptides of the CKP: The peptides are listed from top to bottom in the same order as arrayed in the CKP (N-to C-terminus). The first column of the table contains the names of proteins represented by each peptide and is color-coded to match **Figure 1A**.

### Isotope labeling and purification of the CKP

The yeast strain (HZY3059) carrying the pGal-CKP plasmid (HZE3361) was grown at 30°C in 1 liter of synthetic dropout (-His -Arg) media, supplemented with 30 mg/L light arginine (^12^C_6_ ^14^N_4_) or heavy arginine (^13^C_6_ ^15^N_4_) and 2% raffinose mass/volume (m/v). After cells were grown to saturation, galactose was added to a final concentration of 2% m/v to induce CKP expression for 2 hours (**S2 Figure A**). This produced both a light CKP-^12^C^14^N and heavy CKP-^13^C^15^N, referred to as L-CKP and H-CKP from here onwards (**S2 Figure B**). The cells were then harvested and resuspended in 1.5 mL buffer L (25 mM HEPES-KOH pH 8.0, 175 mM K-glutamate, 2 mM MgCl_2_, 0.1 mM EDTA, 0.5 mM EGTA, 0.1% NP-40, 15% glycerol m/v, protease inhibitor cocktail, and 1 mM PMSF in dH_2_O). This yeast cell resuspension was flash-frozen in liquid nitrogen drop-by-drop (popcorn) and stored at -80 °C. Before the purification, frozen cell popcorns were lysed using a cryogenic grinder (Cat #: 6875D115; SPEX SamplePrep LLC, USA) at 10 cycles per second for 12 cycles, with a 2-minute cycle time and a 2-minute cooling time between cycles. The lysed cell powder was thawed on ice and centrifugated at 21,000 RCF for 20 minutes. 2.5 mL of clarified homogenates were subjected to anti-FLAG-M2 immunoprecipitation with 200 μL of anti-FLAG-M2 agarose resin (Cat #: A2220; Sigma-Aldrich, USA) at room temperature for 1 hour. The resin was washed with 5 mL ice-cold buffer L and 5 mL of 0.1% NP-40. The bound proteins were eluted with 1 mL of 0.1% SDS after 1-hour incubation at 37 °C. The eluate of purified proteins was dried by speed-vac at 55 °C and resuspended in 100 μL of 50 mM ammonium bicarbonate in dH_2_O. To remove detergents, proteins were precipitated using 400 μL of 50% acetone/50% ethanol at -20 °C overnight. Precipitated proteins were centrifuged at 21,000 RCF for 10 minutes, and the resulting protein pellet was washed with 5 mL of 40% acetone/40% ethanol/20% dH_2_O. After washing, the pellet was air dried briefly at 37 °C to evaporate the remaining organic solvent. Trypsin digestion of the protein pellet was performed using 1μg of modified sequencing grade trypsin (Cat #: V5111; Promega, USA) in 100 μL of 20 mM ammonium bicarbonate at 37 °C overnight (12 – 18 hours). Peptides were acidified the following day with 20 μL 10% trifluoroacetic acid (TFA) and diluted to ∼100 fmol/μL as a working stock. Working stock aliquots were stored at -80 °C until use.

### Absolute quantification of the L-CKP and H-CKP via external calibrations

Absolute quantification of CKP was performed using external calibration with a commercially synthesized peptide (GenScript, USA) of GST at its core region (amino acid 104-108: YGVSR). This GST core peptide is expected in the purified CKPs, which share the N-terminal GST tag and show a lack of a noticeable change in electrophoretic mobility (**S2 Figure A**). Thus, this GST core peptide is iso-stoichiometric with CKP peptides. Possible C-terminal degradation of CKP was also excluded since the 3xFLAG tag at its extreme C-terminus was used to purify the full-length CKP. External calibration was performed by preparing 6 calibrators of the GST peptide at 625, 1250, 2500, 5000, 10000, and 20000 pM in 0.1% TFA. Linear regression with 1/x weighting (calibrators with lower concentrations have more weight) was performed using the integrated peak area of each calibrator (**S3 Table**). All calibrator biases were ±15% from target values, and the R^2^ value for the GST peptide was 0.9972. Using this external calibration curve, original stocks of L-CKP and H-CKP were determined to be 140 ± 2 nM and 33 ± 5 nM, respectively (**S3 Figure** and **S3 Table**).

### Validation of absolute quantification using recombinant GST protein

Recombinant GST protein was purified from *E. coli* cells (Rosetta DE3 pLysS Completent Cells, Cat #: 70956; Sigma Aldrich, USA) carrying the plasmid HZE2029 (LIC-2GT) using agarose GS-resin (Cat#: 17-5132-02, GE Healthcare) according to manufacturer instructions. Following purification, GST was precipitated by 50% acetone/50% ethanol and solubilized in 6M urea/50 mM ammonium bicarbonate in deionized water (dH_2_O). The concentration of GST was quantified using Beer’s Law (Molar extinction coefficient for GST: ε = 44,350 cm^-1^ M^-1^) and absorption at 280 nm on a NanoDrop^TM^ spectrophotometer (Cat #: ND2000; Thermo Scientific, USA). The GST stock concentration was determined to be 0.91 ± 0.01 μM (**S4 Figure**). This stock was then diluted to 2.5 nM, subjected to trypsin digestion, and quantified using the external calibration curve of the commercially synthesized GST peptide to validate the external calibration curve.

### Reverse-phase Liquid chromatography

The liquid chromatography method utilized two solvents: Mobile phase A (MPA, 0.1% formic acid in dH_2_O) and mobile phase B (MPB, 0.1% formic acid in ACN). Chromatography was obtained using a Vanquish Neo UHPLC with a self-packed fused silica (Cat #: 2000023; Polymicro, USA) C18 column (170 mm length × 100 μm I.D; 2.2 μm particle size, Cat#: 101182-0000, Sepax, USA). Samples were injected using a backward flush Heated Trap-and-Elute workflow (50 °C) on a PepMap Neo Trap cartridge (Cat#: 174500; Thermo Fisher, USA). The analytical column was kept at 50°C using a Sonation PRSO-V2 (Sonation GmbH, Germany) column oven. The flow rate was 2.25 μL/min across the entire method, and the initial starting conditions were 98% MPA and 2% MPB. The linear gradient begins at 5.0 min, at which point the composition of MPB increases linearly to 40% across 45.0 min. At 50.0 min, MPB was increased linearly to 95% over 5.0 min, where it was held constant for 5.0 min before returning to 2% MPB over 0.1 min. Equilibration was performed for the final 5.0 min at 2% MPB. The total LC method is 65.1 min and the injection volume for all analyses was 3 μL.

### Parallel-Reaction Monitoring Mass Spectrometry (PRM-MS)

MS analysis was performed in positive ion mode on a Q-Exactive Plus mass spectrometer (Thermo Fisher, USA) using a Nanospray Flex Ion Source (Cat #: ES071; Thermo Fisher, USA). Parallel-Reaction-Monitoring (PRM) was performed in data-independent acquisition mode with a targeted inclusion list (**S4 Table**) and a single Full MS scan accompanying each set of PRM scans. Source parameters are as follows: Spray voltage – 2.75 kV, Capillary temperature – 290 °C, S-lens RF level – 50.0. Full MS parameters are as follows: Resolution – 35,000, AGC Target – 1e6, Maximum IT – 50 ms, and Scan range – 280 – 900 m/z. PRM-MS parameters were: Resolution – 17,500, AGC Target – 5e5, Maximum IT – 100 ms. Data was acquired in Xcalibur (Version 4.5.474.0) and analyzed in Skyline (Version 23.1) [61,62].

### Evaluation of peptide collision energy

The effect of collision energy on product ion distribution and signal intensities were evaluated by performing replicate injections of ∼50 fmol of H-CKP. Across these injections, the collision energy (CE) and normalized collision energy (NCE) were raised by increments of 2% from 10% (lower bound of the instrument) until a Gaussian distribution of chromatographic peak areas were observed for the product ion breakdown curves of each peptide. Integrated peak areas for each peptide’s top 10 visible transitions were calculated in Skyline and exported into Excel for subsequent analysis. Optimal collision energies were determined for each peptide based on overall product ion intensities. The collision energies used for the PRM-MS method (**S4 Table**) were selected with additional considerations of matrix interferences and quantifier ion signal intensities.

### Generation of stabilization matrix

The matrix used in this study was derived from a tryptic digest of yeast cytosolic proteins obtained through chromatin fractionation, as previously described [63]. Briefly, 200 OD_600_ · mL of yeast cells were treated with Lyticase (in-house purified, 0.5 mg/mL, 800 μL) to generate spheroplasts [64]. Spheroplasts were lysed with 0.5% Triton X-100 on ice for 20 minutes, and the lysate was spun through a 30% sucrose cushion to separate the cytosolic proteins from the chromatin. The soluble fraction (including the sucrose layer) was isolated, reduced with 10 mM DTT at 37 °C for 30 minutes, alkylated with 30 mM iodoacetamide at room temperature in the dark for 15 minutes, precipitated by 50% ethanol/ 50% acetone, and digested into peptides by trypsin. The total protein concentration of the matrix was quantified by Bradford assay (BioRad, USA). A working stock of the matrix peptides, referred to as matrix here onwards, was prepared at 50 ng/μL in 0.1% TFA and stored at -80 °C until used.

### Generation of isotope dilution external calibration curves

Working stocks of calibrators (L-CKP) and the internal standard (H-CKP) were prepared in 50 ng/μL matrix at 120 nM and 2 nM following absolute quantification. The L-CKP was serially diluted to generate a total of 10 working stocks with concentrations ranging from 156.3 pM to 120 nM, one for each calibrator level. Calibrators were prepared in glass inserts (Cat#110000101; DWK, GmbH) by mixing 20 μL of the working stocks of L-CKP with 20 μL of the working stock of H-CKP (40 μL final volume). The final concentration of H-CKP (internal standard) in-vial was 1 nM for each of the 10 calibrators, and the final concentration of L-CKP ranged from 78.1 pM to 60 nM. The resulting peak area ratios between L and H-CKP were plotted as a function of L-CKP concentrations in Skyline. Calibration curves were generated by fitting a linear regression with 1/x weighting (calibrators at lower concentrations have more weight) to the plot. Calibrators were excluded if they failed any of the following criteria: 1) signal to noise ratio greater than 10, 2) accuracy (%Bias) within ± 20% of target calibrator concentration, and 3) imprecision (%CV) of less than 15% across replicate analysis. Calibration curves generated were only used for downstream quantification if they had an R^2^ (coefficient of determination) > 0.99 and were fitted to at least 4 calibrators (had a minimum of 4 points on the curve).

### *Ex vivo* reconstitution of budding yeast kinetochores

#### Yeast cell extract preparation

Yeast cells (HZY1028) were grown in 1 liter of YPD (yeast extract, peptone, dextrose; Fisher Scientific, USA) medium at 30 °C to OD_600_ ∼ 0.3 and arrested in M or G1-phase by adding Nocodazole to a final concentration of 150 μg/mL or alpha factor to a final concentration of 15 nM, respectively, for 3 hours. After the arrest, cells were centrifuged at 4,000 RCF, and the resulting cell pellet was washed once with phosphate-buffered saline (PBS, pH 8.0) and resuspended in 1/4 cell pellet volume buffer L. The resuspended cell slurry was flash-frozen in liquid nitrogen and lysed by a Cryogenic Grinder (Cat #: 6875D115; SPEX SamplePrep LLC, USA) as described above. The protein concentration of the clarified yeast extract was ∼ 80 mg/mL by Bradford assay (BioRad, USA). The extract was stored at - 80 °C until reconstitution experiments were performed.

#### Preparation of centromeric and control DNA beads

Yeast centromere III (CEN DNA), centromere III with point mutations to its CDE III region (MUT DNA), and yeast autonomously replicating sequence (ARS DNA) were amplified from HZE3240, HZE3241, and HZE3246, respectively, under standard PCR conditions with 0.5 μg Taq polymerase (in-house purified)/20 mM Tris-HCl (pH 8.4)/50 mM KCl/dNTP (50 μM each)/2 mM MgCl_2_ in dH_2_O and primers 1 and 2 for CEN and MUT DNA, and primers 3 and 4 for ARS DNA: 1) CEN3-For: 5’ – GGCGATCAGCGCCAAACA – 3’; 2) Bio-CEN3-Rev: 5’ - /5Biosg/CGCTCGAATTCGGATCCG – 3’; 3) TRP1-For: 5’ – GAAGCAGGTGGGACAGGT – 3’; and 4) Bio-ARS-Rev: 5’ -/5Biosg/CCCCCTGCGATGTATATTTTC – 3’. To improve PCR efficiency, additional dTTP and dATP were supplemented (to a final concentration of 200 μM each) when amplifying the AT-rich CEN and MUT DNA. The resulting PCR product was 411 bps in length. PCR products were precipitated and washed with 70% ethanol/30%100mM ammonium acetate in dH_2_O. The DNA pellet was resuspended in Tris-EDTA buffer (pH 7.5), and the concentrations of the purified DNA were measured by a NanoDrop^TM^ spectrophotometer (Cat #: ND2000; Thermo Scientific). Before each reconstitution, 50 μL of Dynabeads^TM^ M-280 streptavidin (Cat #: 11206D; Thermo Scientific) were incubated with 2-3 μg of the purified DNA at room temperature for 20 minutes in 1M NaCl/0.5nM EDTA/5mM Tris-HCl (pH 7.5) in dH_2_O. The amount of DNA bound to the beads was calculated to be ∼1.5 μg per 100 μL of the original bead solution. After binding with DNA, the beads were washed 2 times by buffer L and stored at 4 °C until use.

#### Kinetochore reconstitution

Frozen yeast extracts were thawed from -80 °C on ice before reconstitution experiments. For each kinetochore reconstitution, 50 μL of DNA beads were incubated with 500 μL of yeast extract at room temperature for 1 hour. Following incubation, the beads were washed 4 times with 500 μL ice-cold buffer L. Each wash was 500 μL and the last 2 washes were allowed to incubate with the beads at room temperature for 2 minutes before buffer removal. To elute reconstituted kinetochores, the washed beads were resuspended in 100 μL buffer L with 0.1 U/μL Turbo DNase-I (Cat #: AM2238, Thermo Fisher Scientific) and incubated at room temperature for 1 hour. Eluates were reduced, alkylated, and precipitated by 50% ethanol/ 50% acetone and prepared for MS analysis as described above. Normalized concentrations of kinetochore proteins were calculated by dividing the in-vial concentration, as determined by LC-PRM-MS, by the percent of total sample injected (e.g. 3 nM / 7.5% = 40 nM; 3 nM is the measured in-vial concentration and 40 nM is the normalized concentration, 7.5% is the amount of total sample injected for LC-PRM-MS).

## Results

### Design of the concatenated kinetochore protein (CKP)

The CKP contains 25 concatenated tryptic peptides derived from kinetochore subunits (**Figure 1A and 1B**), whose sequences are listed in **Table 1**. CKP expression was performed in yeast using a high-copy expression plasmid with a strong galactose-inducible promoter to maximize yield. C-terminal arginine peptides were selected to facilitate the incorporation of isotopically light (^12^C_6_ ^14^N_4_) or heavy arginine (^13^C_6_ ^15^N_4_) via the SILAC approach in an arginine auxotrophic yeast strain [65,66]. To promote robust protein expression, stability, and ease of purification, the CKP is flanked by N-terminal GST and C-terminal 6xHIS-3xFLAG tags. The N-terminal GST, a globular protein, aids with CKP solubility during its expression, while the C-terminal tags were used to purify the CKP. Expression and purification of both isotopically light CKP (L-CKP) and isotopically heavy CKP (H-CKP) were evaluated by anti-FLAG Western blotting or Coomassie staining (**S2 Figure**). The Western blot and Coomassie staining confirmed that full-length CKPs were purified, ensuring that tryptic peptides of the kinetochore are iso-stoichiometric within the CKPs.

**Figure 1.**
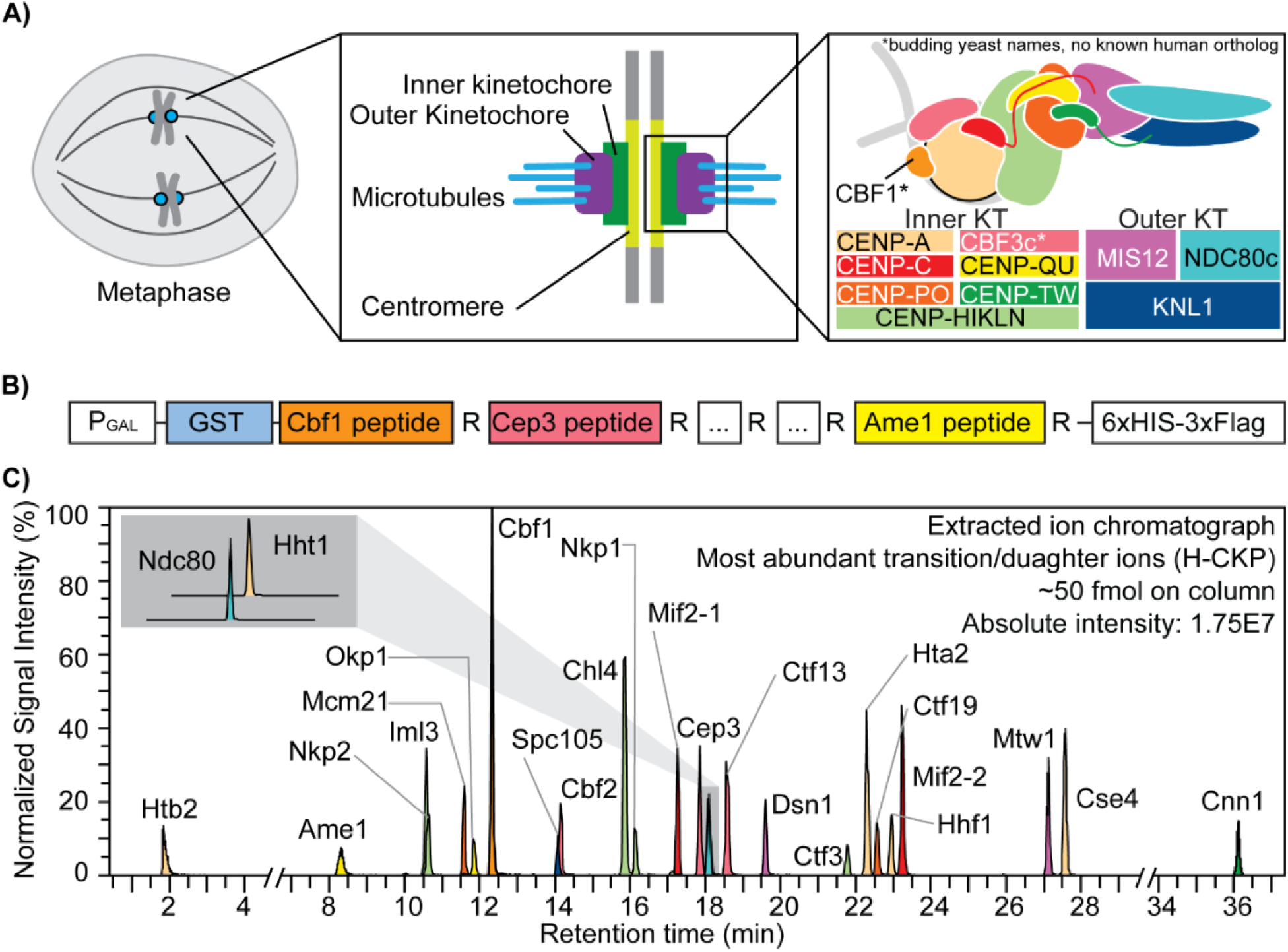
The kinetochore and the concatenated kinetochore protein (CKP) construct. **A)** Schematics of yeast kinetochore organization: Yeast kinetochores assemble on the point centromere at each chromosome to carry out mitotic chromosome segregation. The kinetochore can be divided into inner and outer sub-complexes, where the inner kinetochore contacts the centromeric chromatin, and the outer kinetochore forms an association with the spindle microtubules. Both inner and outer kinetochores are composed of multiple protein complexes, as illustrated and annotated in the right-most panel. **B)** A gene block is constructed to express and purify the concatenated kinetochore protein (CKP): 25 tryptic peptides with a C-terminal arginine were selected from DDA-MS results and incorporated into the CKP. The expression of the protein is controlled by a galactose-inducible promoter (P_GAL_) in yeast. **C)** Extracted ion chromatograph of the most abundant daughter ions of H-CKP peptides with retention time in minutes. The y-axis indicates the signal intensities of each ion expressed as a percentage of the signal from the ion with the highest signal intensity (Cbf1 peptide). The total amount of H-CKP injected and the absolute signal intensity for the Cbf1 peptide’s most abundant product ion are labeled on the top right of the plot.

Next, PRM scans of both light and heavy tryptic peptides of the CKP were incorporated into an initial targeted LC-MS/MS method using a normalized collision energy (NCE) of 27%. Using this PRM method, extracted product ion chromatographs of H-CKP peptides (**Figure 1C**) confirmed the thorough incorporation of the heavy stable isotopes (> 99%) with undetectable signal observed for PRM scans of the corresponding light peptide (**S5 Table**). Detection sensitivity by PRM-MS differs according to peptide sequence and corresponding chemical properties, and this produces varying observed product ion intensities (**Figure 1C**).

### Evaluation of peptide collision energy

Collision energy (CE) was evaluated for all 25 peptides of the CKP (**Figure 2**) on column by replicate injections. Collision energy (CE) or normalized collision energy (NCE) was increased in series by 2% for each injection until product ion signal intensities decreased. The resulting breakdown curves for two representative peptides are shown in **Figures 2A** and **2B**. Gaussian distributions were observed for the Mcm21 peptide (IDDISTSDR) and the Nkp1 peptide (EIYDNESELR) between 10% and 22% CE (**Figure 2A, 2B**). Optimal CEs of all peptides were determined based on the overall signal intensities of the top product ions. CE fragmentation produced greater changes in product ion signal intensity relative to NCE fragmentation, and the highest product ion signal intensities were observed when utilizing the CE fragmentation scheme for all peptides but the Htb2 peptide (**Table 2**).

**Figure 2.**
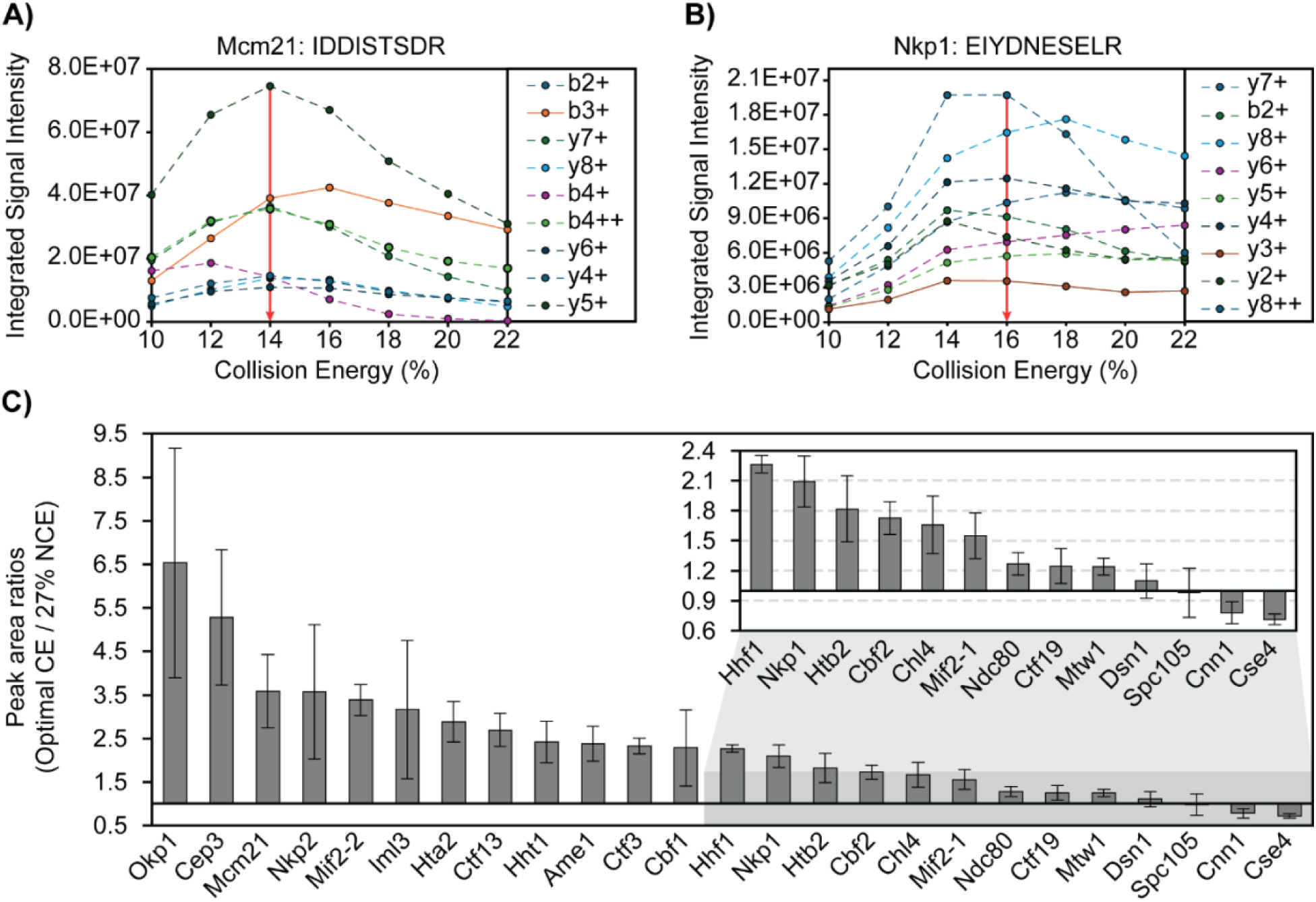
Collision energy is optimized for each CKP peptide to improve signal intensity and transition heterogeneity. **A & B)** Representative breakdown curves of IDDISTSDR (Mcm21 peptide) and EIYDNESELR (Nkp1 peptide): The peak areas of the top 9 most abundant product ions were plotted against %CE to illustrate the changes in fragmentation patterns of the peptide in response to changes in collision energies. **C)** Peak area ratios of product ions before and after CE optimization: The mean peak area of each peptide’s most abundant product ions (sum of peak areas of the top 10 most abundant product ions) across 3 analytical replicates were plotted for injections using either 27% NCE for all peptides or optimal CEs (see Table 2). Mean peak area ratios were plotted in descending order as grey bars and standard deviations were plotted as error bars. The x-axis intersects the y-axis at y = 1, which indicates no improvement of product ion signal intensity from CE optimization.

**Table 2.** Summary of optimal collision energies: Parentheses indicate NCE was used. Otherwise, CE was used.

| **Protein** | **Peptide** | **(N)CE** |
| --- | --- | --- |
| Cbf1 | LSTEDEEIH SAR | 26 |
| Cep3 | LVYLTER | 10 |
| Ctf13 | TGLADFTR | 12 |
| Cse4 | YTPSELALYEIR | 19 |
| Htb2 | HAVSEGTR | (20) |
| Hta2 | AGLTFPVGR | 10 |
| Hhf1 | ISGLIYEEVR | 16 |
| Hht1 | STELLIR | 10 |
| Mif2-1 | YSLDTSESPSVR | 22 |
| Mif2-2 | VAPLQYWR | 12 |
| Cbf2 | EENIVNEDGPNTSR | 25 |
| Mcm21 | IDDISTSDR | 14 |
| Ctf19 | QQLSLLDDDQVR | 24 |
| Ctf3 | DAPGSATLILQR | 16 |
| Iml3 | ESIVTSTR | 10 |
| Chl4 | NEDSGEPVYISR | 20 |
| Mtw1 | IPEEYLDANVFR | 25 |
| Cnn1 | SFLQDLSQVLAR | 14 |
| Nkp1 | EIYDNESELR | 16 |
| Nkp2 | VTSELEAR | 10 |
| Ndc80 | QYDSSIQNLTR | 23 |
| Dsn1 | ILDNTENYDDTEL R | 28 |
| Spc105 | VHISTQQDYSPSR | 10 |
| Okp1 | VIQAEYR | 10 |
| Ame1 | NDEDLTTR | 12 |

Signal intensities of each peptide’s top 10 most abundant ions were compared before and after CE optimization (**Figure 2C**). Optimal CEs were compared to NCE at 27%, a commonly used CE for proteomics analysis on similar instrumentation [50]. Improved signal intensities were observed for 21 peptides, and no improvements were observed for the Dsn1, Spc105, Cnn1, and Cse4 peptides. (**Figure 2C**). The relative ion intensity among product ions (ion ratio) is unique to a peptide of a defined amino acid sequence at a given collision energy. Thus, having diverse product ions allows for the selection of different product ions as quantifiers for quantification and qualifiers to confirm identification and resolve interferences. PRM-MS monitored all daughter ions during the evaluation of CE and NCE, which allowed us to revise the optimal NCE/CEs after taking into consideration matrix interferences and quantifier ion signal intensities for the final PRM-MS method (**S4 Table**).

### Peptide stability and matrix stabilization of peptides

Due to the ∼1 hour duration of the LC-PRM-MS method, an analytical batch can take over 24 hours to complete, and peptide digests are kept at 10 °C in the autosampler for this duration. Thus, peptide stability was evaluated by analyzing replicate injections (n=6) of digested H-CKP kept at 10 °C in the autosampler across a period of ∼30 hours, and recovery (peak area) for all 25 peptides reconstituted in 0.1% TFA (no matrix) were monitored (**Figure 3**). After 15 hours, 18 peptides were recovered > 70% of initial levels, 2 peptides between 70% and 50%, and 5 peptides < 50% (**Figure 3C** and **Table 3**). At 30 hours, 6 peptides were recovered > 70% of initial levels, 10 peptides between 70% and 50%, and 9 peptides < 50% (**Figure 3C** and **Table 3**). Thus, most peptides had poor recoveries across this period, indicating poor stability at 10 °C. Some peptides had particularly poor recoveries, which include those derived from Cnn1, Cse4, and Mif2. Common features of these peptides include that they elute at later retention times on the LC gradient (**Figure 1C**) and contain a higher proportion of large hydrophobic amino acid residues such as tryptophan, phenylalanine, leucine, and isoleucine relative to other targeted peptides (see **Table 1** for peptide sequences).

**Figure 3.**
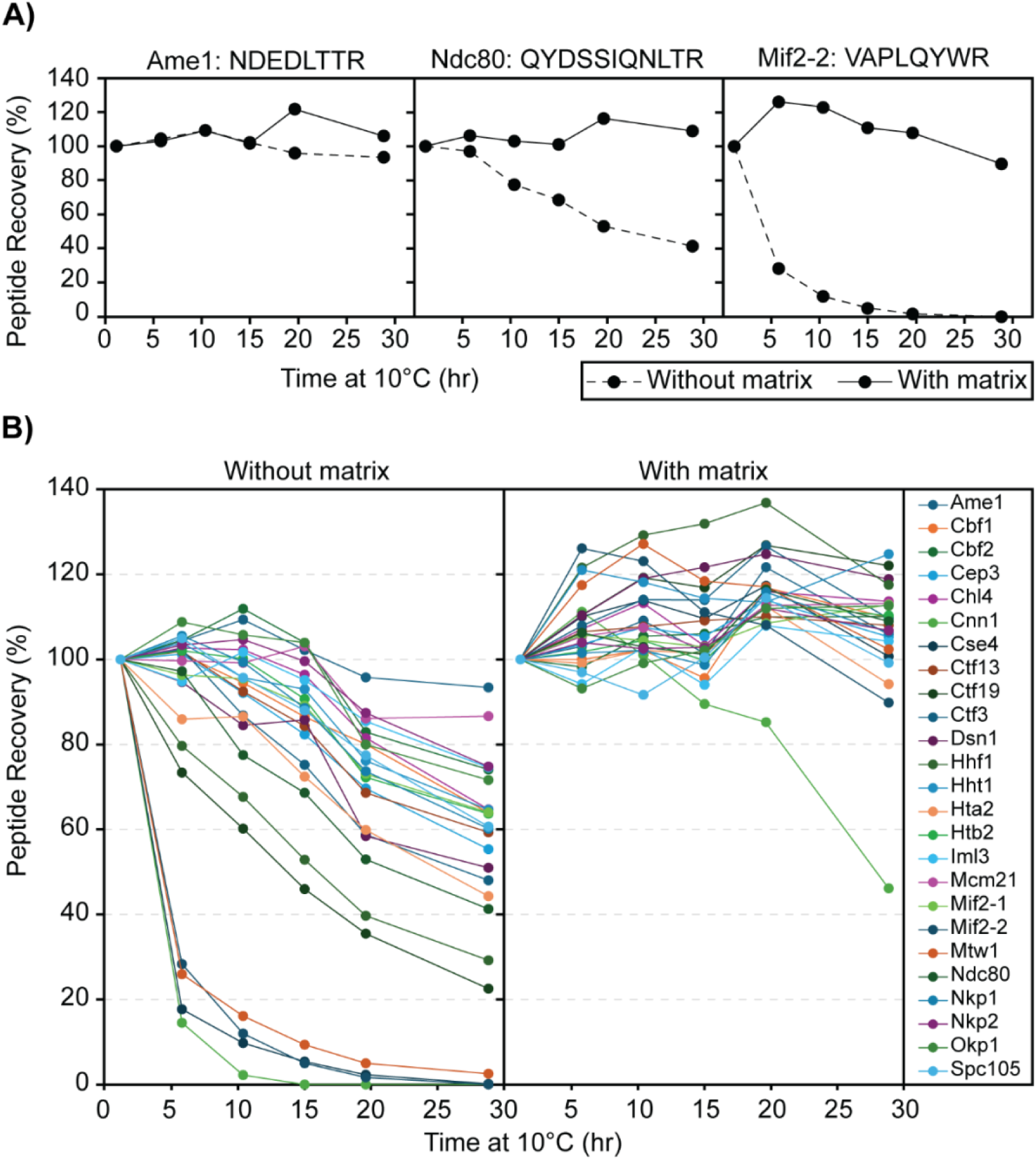
Effects of stabilization matrix on peptide recovery. **A)** Representative peptide recovery profiles over time: No matrix (0.1% TFA) - solid line; With matrix (0.1% TFA with 50 ng/μL digested yeast cytosolic proteins) - dashed line. The peptides’ sequences are labeled on the top of each panel. The y-axis shows the percentage recovery of the peptide by peak area in comparison to the first time point (1 hour at 10°C), and the x-axis shows the amount of time that the digested CKP was left at 10°C before injection into the MS. **B)** Recovery profiles of all CKP peptides: As above, the percentage recovery of each peptide was plotted against time. Each peptide is marked by a different color, as indicated by the legend key on the right-most panel.

**Table 3.**
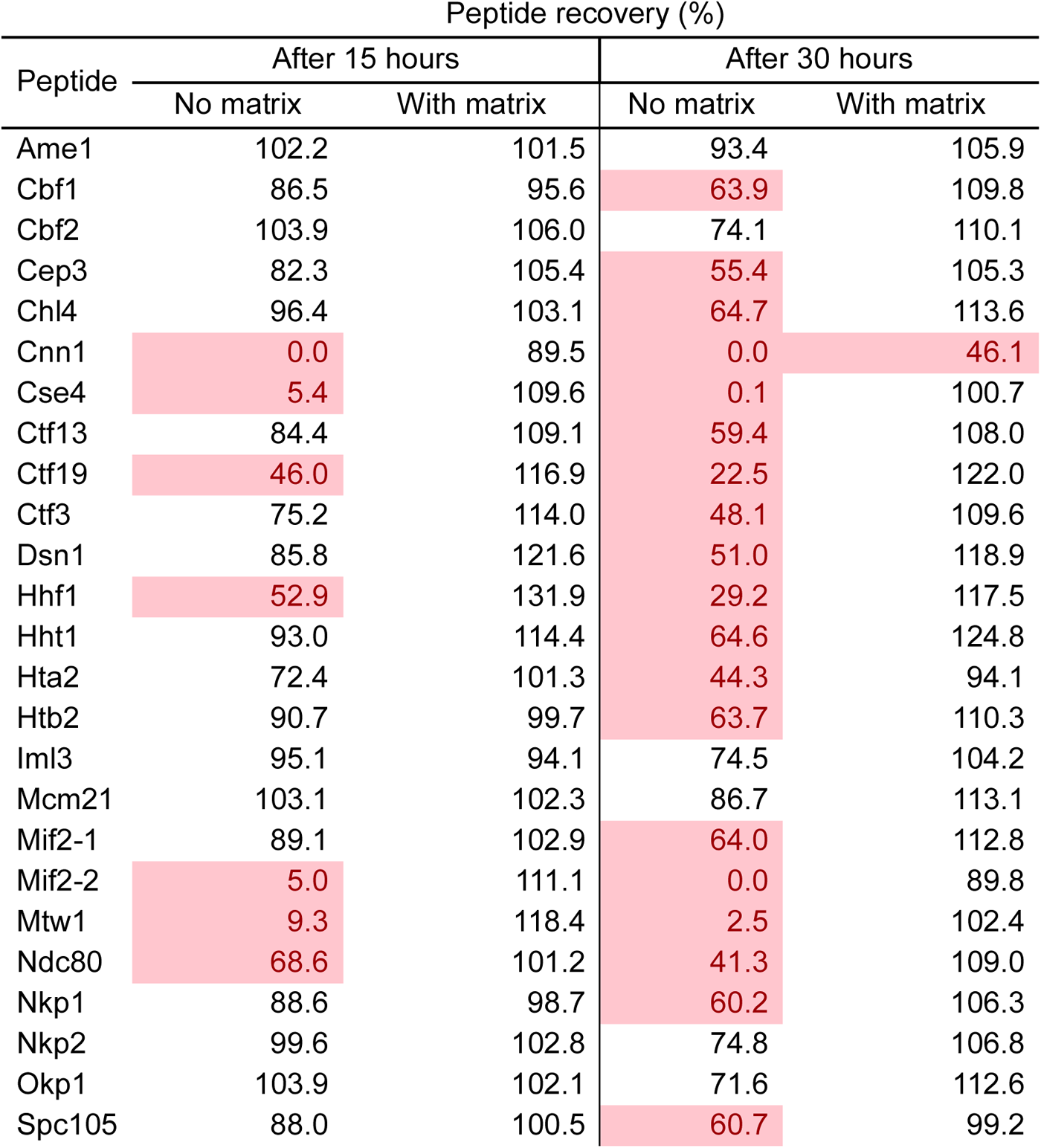
Summary of Peptide Recovery Pre- and Post-Matrix Matching: Red text with red highlights indicates peptide recovery < 70%.

To address the poor peptide recovery, 50 ng/μL of digested yeast cytoplasmic proteins was included as a stabilizing matrix to circumvent losses from peptide precipitation or non-specific adsorption losses in-vial. The effect of matrix stabilization was evaluated, as shown in **Figure 3**, following the reconstitution of digested H-CKP in the stabilization matrix instead of 0.1% TFA. After 30 hours in a 50 ng/μL stabilization matrix, all peptides were recovered at > 85% of initial levels except for the Cnn1 peptide. Recovery for the Cnn1 peptide was 89.5% at 15 hours and 46.1% at 30 hours (**Table 3**). Direct comparisons of the matrix effects on peptide recoveries are illustrated in **Figure 3A** for 3 representative peptides of varying recoveries across 30 hours. Thus, the stabilization matrix significantly improved the recoveries of all 25 peptides over a 30-hour period (**Figure 3B**). The matrix was necessary for maintaining peptide stability and facilitating acceptable recoveries during subsequent method validation and data analysis.

### Evaluation of the analytical measurement range and limits of quantification

Evaluation and establishment of the analytical measurement range (AMR) were performed to ensure robust peptide quantification. To determine the AMR for each peptide, including the lower limit of quantification, 10 calibrators comprised of digested L-CKP with concentrations ranging from 78 pM to 60 nM in-vial were mixed with a fixed concentration of internal standard (1 nM of H-CKP) in-vial and reconstituted in the stabilization matrix. Utilizing the peak-area-ratio (analyte peak area/internal standard peak area), external calibration curves for all 25 peptides were generated with the requirement of a minimum of four calibrator levels, calibrator biases within ± 20% from nominal target values (**S6 Table**), and R^2^ values greater than 0.99 (**Table 4**).

**Table 4.** Summary of CKP Peptide Measurement Linearity and Limits of Quantification. LLoQ: lower limit of quantification, ULoQ: upper limit of quantification.

| Range | Peptide | R<sup>2</sup> | Slope | Intercept | # of Calibrators | LLOQ (pM) | ULoQ (pM) |
| --- | --- | --- | --- | --- | --- | --- | --- |
| Low | Hta2 | 0.9998 | 1.03E-03 | 8.89E-03 | 9 | 78 | 60000 |
| Low | Ctf3 | 0.9999 | 1.89E-03 | -4.49E-02 | 9 | 78 | 60000 |
| Low | Cbf2 | 0.9964 | 9.59E-04 | 1.55E-01 | 7 | 625 | 60000 |
| Low | Nkp1 | 0.9966 | 1.03E-03 | 7.86E-02 | 7 | 625 | 60000 |
| Low | Iml3 | 0.9991 | 9.25E-04 | 5.82E-03 | 10 | 78 | 60000 |
| Low | Htb2 | 0.9999 | 8.34E-04 | 1.12E-01 | 7 | 625 | 60000 |
| Low | Mcm21 | 0.9997 | 1.03E-03 | 4.15E-03 | 10 | 78 | 60000 |
| Low | Dsn1 | 0.9999 | 1.12E-03 | -2.81E-02 | 9 | 78 | 60000 |
| Low | Mtw1 | 0.9935 | 8.03E-04 | -1.66E-01 | 4 | 313 | 2500 |
| High | Mtw1 | 1 | 1.39E-03 | -1.55E+00 | 4 | 2500 | 20000 |
| Low | Hhf1 | 0.9996 | 7.49E-04 | -2.86E-01 | 5 | 1250 | 60000 |
| Low | Cbf1 | 0.9971 | 8.05E-04 | 3.77E-03 | 9 | 78 | 60000 |
| Low | Cep3 | 0.9952 | 6.99E-04 | 6.53E-03 | 10 | 78 | 60000 |
| Low | Ame1 | 0.9987 | 1.12E-03 | -3.23E-02 | 10 | 78 | 60000 |
| Low | Chl4 | 0.9996 | 9.41E-04 | 2.32E-02 | 10 | 78 | 60000 |
| Low | Ctf19 | 0.9998 | 1.48E-03 | -8.24E-02 | 8 | 78 | 60000 |
| Low | Ndc80 | 0.9998 | 1.12E-03 | -4.17E-03 | 9 | 78 | 60000 |
| Low | Cnn1 | 0.9903 | 7.32E-04 | -1.44E+00 | 4 | 2500 | 20000 |
| Low | Hht1 | 0.9999 | 8.06E-04 | 1.39E-01 | 8 | 313 | 60000 |
| Low | Ctf13 | 0.9995 | 7.87E-04 | 2.12E-02 | 8 | 156 | 20000 |
| Low | Mif2-2 | 0.9992 | 1.04E-03 | -2.91E-02 | 8 | 78 | 60000 |
| Low | Spc105 | 0.9999 | 2.43E-03 | -3.13E-02 | 7 | 313 | 60000 |
| Low | Okp1 | 0.9993 | 7.33E-04 | 2.99E-02 | 9 | 156 | 60000 |
| Low | Nkp2 | 0.9998 | 7.94E-04 | 5.98E-02 | 8 | 313 | 60000 |
| Low | Mif2-1 | 0.9997 | 8.21E-04 | 1.57E-01 | 6 | 625 | 60000 |
| Low | Cse4 | 0.9912 | 4.14E-04 | -7.55E-02 | 4 | 313 | 2500 |
| High | Cse4 | 0.9988 | 8.32E-04 | -1.28E+00 | 5 | 2500 | 60000 |

All calibrators were evaluated for inter-day accuracy and precision through analytical replicates (**S7 Table** for inter-day accuracy and **S8 Table** for inter-day precision). Product ions were chosen for peptide quantification (quantifier ions) and identification (qualifier ions) based on signal intensity and absence of matrix interference (**S9 Table** for L-CKP ion masses and **S10 Table** for H-CKP ion masses).

The lower limit of quantification for each peptide was determined as the lowest calibrator level with signal-to-noise ratio greater than 10 in the quantifier ion, average biases within ± 20% for accuracy, and %CVs < 15% for precision. The AMR for each peptide was then established as the range from the lower limit of quantification up to the highest calibrator level which met accuracy and precision acceptability criteria. In summary, most peptides (84%) have AMRs which span across two orders of magnitude (**Figure 4**). **Figure 4A** shows an example peptide with an AMR spanning 10 calibrators, and **Figure 4B** shows an example peptide with a much more limited AMR. **Figure 4C** and **Table 4** summarize the AMRs of all CKP peptides. The Mtw1 and Cse4 peptides have 2 AMRs, which require separate calibration curves (as shown in **Figure 4C** by the 2 horizontal bars with different shades of grey). **S5 Figure** shows the high and low-range calibration curves of Cse4 as an example.

**Figure 4.**
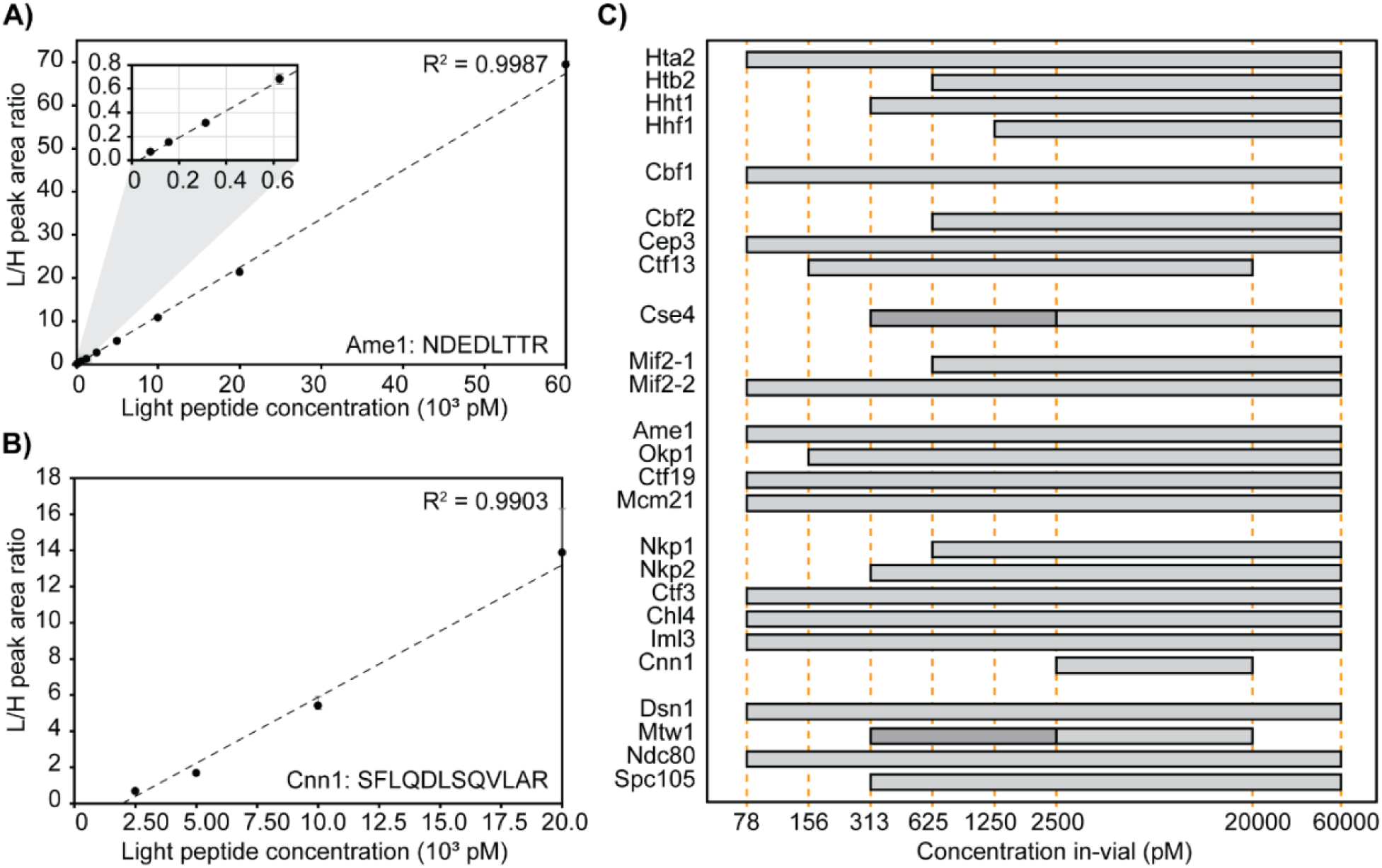
Linearity and limits of quantification. **A & B)** Representative calibration curve of the Ame1 peptide (NDEDLTTR) and the Cnn1 peptide (SFLQDLSQVLAR). In-vial concentrations (pM) are shown on the x-axis, and the peak area ratio between L-CKP and H-CKP is plotted on the y-axis. A zoom region inset of the plot is appended in the top left corner to show the curve fitting at the lower calibrators. **C)** AMRs of all CKP peptides: The AMRs of CKP peptides are plotted as grey bars on a log_10_ scale. Orange dashed lines intersecting the x-axis indicate calibrator concentrations (pM). Dark grey bars indicate the lower AMRs of Cse4 and Mtw1 peptides.

### CASA of *ex vivo* reconstituted kinetochores using M-phase yeast cell extracts

To demonstrate the suitability of this LC-PRM-MS method for determining protein complex stoichiometry, *ex vivo* reconstituted kinetochores were analyzed (**Figure 5**). *Ex vivo* yeast kinetochores were reconstituted on biotinylated DNA-bound streptavidin beads using yeast whole-cell extracts derived from cells arrested in M-phase by nocodazole (**Figure 5A**). Reconstituted kinetochores were eluted from DNA beads using DNase-I, and eluates were prepared for tryptic digestion and subsequent isotope dilution LC-PRM-MS analysis. The reconstitution and DNase I elution efficiency was first evaluated by Western blotting using a Mif2-TAF tagged strain (HZY2347) (**S6 Figure**). After this, a *bar1Δ* (HZY1029) was used for all subsequent reconstitutions for PRM-MS. Kinetochore reconstitution on three different types of DNA beads were evaluated using process replicates (n=3): CEN-DNA (wild-type yeast centromere III), MUT-DNA (yeast centromere III with a point mutation at CDE-III), and ARS-DNA (autonomously replicating sequence) (**Figure 5A**).

**Figure 5.**
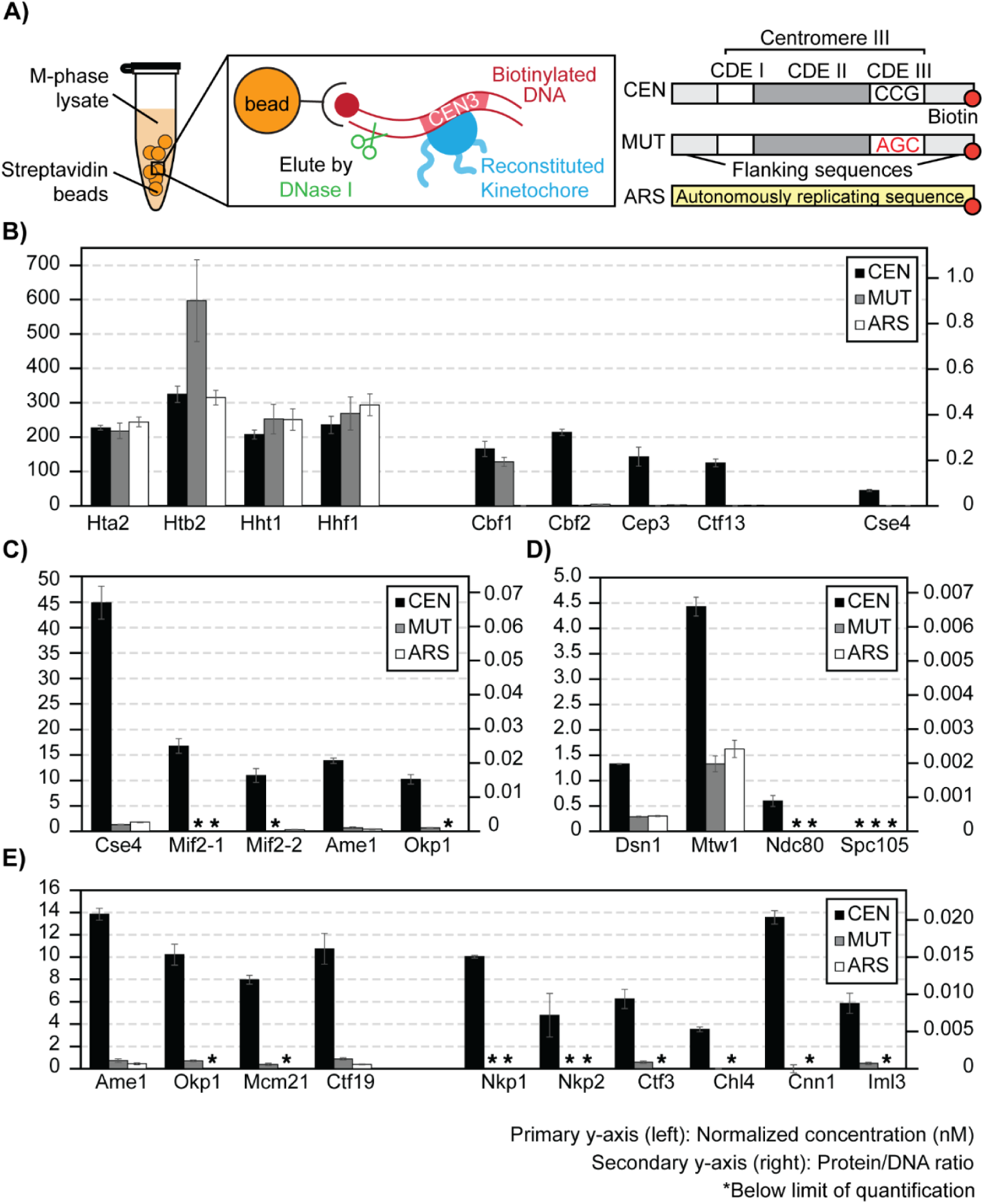
Stoichiometry of M-phase yeast kinetochores. **A)** Right: schematics of *ex vivo* kinetochore reconstitution using yeast whole cell extracts (WCEs); Left: DNA maps for the 3 types of DNA used. **B)** Quantification of *ex vivo* reconstituted nucleosomes, Cbf1, and the Cbf3 complex: The primary y-axis (left) shows the normalized concentration of proteins (see method). The secondary y-axis (right) shows the amount of each protein as a ratio to the total amount of DNA on beads. The standard deviation of 3 process replicates (3 *ex vivo* reconstitutions done in parallel) are plotted as error bars around their averages. Quantifications are only made when the measurement falls within the AMRs defined in Figure 4. * Indicates signals below the lower limit of quantification. **C)** Quantifications of *ex vivo* reconstituted essential yeast inner kinetochore proteins: Cse4, Mif2, Ame1, and Okp1. **D)** Quantifications of *ex vivo* reconstituted outer kinetochore subunits: Dsn1, Mtw1, Ndc80, Spc105. **E)** Quantifications of *ex vivo* reconstituted COMA complex and Ctf3 complex members.

Canonical histones associated readily with DNA, regardless of sequence. All four histone subunits were determined to be iso-stoichiometric in all studies, except for fluctuations in the amount of Htb2 associated with MUT-DNA (**Figure 5B**). This Htb2 peptide suffers from significant ion suppression in the reconstitution sample matrices, leading to low internal standard recovery (**S7 Figure**). Cbf1 was robustly reconstituted on CEN-DNA and MUT-DNA but not on the ARS-DNA (200-fold enrichment, **Figure 5B**). Indeed, Cbf1 binds the E-box consensus sequence (CACGTG) in the CDE-I region of the centromere [67]. The CBF3 complex (Cbf2-Cep3-Ctf13-Skp1; Skp1 was not monitored here) bound specifically to the CEN-DNA and not to the MUT-DNA or the ARS-DNA (**Figure 5B**). This confirms the specific role of the CBF3 complex in recognizing the conserved CCG element of CDE-III [41,68], which was mutated to AGC in the MUT-DNA. The three monitored subunits of the CBF3 complex have an average calculated integer stoichiometry of 12: 8: 7 (Cbf2: Cep3: Ctf13), which deviates from the stoichiometry of *in vitro* reconstituted CBF3 complexes (2: 2: 1; Cbf2: Cep3: Ctf13) [20,22]. However, a stoichiometry of 2: 2: 1 (Cbf2: Cep3: Ctf13) can be assigned to the complex while staying within the 95% confidence intervals of the peptide measurement (n=3).

The centromere-specific H3 variant, Cse4, was observed to reconstitute specifically on the CEN DNA (∼ 20-fold higher) relative to the MUT or ARS DNA (**Figure 5B**). Unlike histone H3, Hht1, Cse4 loading is much less efficient (∼ 4-fold less). The amount of Cse4 on the CEN-DNA was approximately 4-fold lower than Cbf2. Cse4 is expected to 1 : 1 to Cbf2 when completely loaded [23], indicating that not all loaded CBF3 complexes was associated with Cse4. Likewise, the observed Cse4 : Mif2 stoichiometry was approximately 3 : 1, indicating that not all Cse4 nucleosomes were associated with Mif2 (**Figure 5C**). Notably, the two peptides of Mif2 did not quantify iso-stoichiometrically, showing a deviation of 34% (**Figure 5C**). Analytical variability was ruled out since both Mif2 peptides were quantified reproducibly within the AMR with %CVs < 15% from three process replicates (**S11 Table**). In addition, the calculated average integer stoichiometry of Ame1 : Okp1 was 4 : 3, while 1 : 1 stoichiometry is expected as Ame1 and Okp1 form a stable heterodimer [69]. However, 1 : 1 stoichiometry (Ame1: Okp1) can be assigned while staying within their 95% confidence intervals. Though modest, such discrepancies should be addressable with additional Mif2, Ame1, and Okp1 peptides (**See Discussion**). Interestingly, the stoichiometry between Mif2 and Ame1-Okp1 was close to 1:1, indicating that these proteins reconstituted at a comparable level on the Cse4-bound CEN-DNA (**Figure 5C**). The outer kinetochore subunits reconstituted on centromere DNA at further reduced levels relative to the inner kinetochore subunits (**Figure 5D)**. Finally, the remaining non-essential inner kinetochore subunits were quantified at reduced levels relative to Mif2 and Ame1-Okp1 (**Figure 5E**).

Altogether, under the assumption that each molecule of DNA associates with one centromeric or canonical nucleosome, we calculated that ∼40% of the CEN-DNA is non-specifically associated with canonical nucleosomes, ∼25% is associated with Cbf1 and the Cbf3 complex, but only ∼7% is associated with Cse4. Thus, Cse4 recruitment is a critical limiting factor of this *ex vivo* kinetochore assembly system. Subsequent assembly is not stoichiometric, as only ∼2% of CEN-DNA is associated with Mif2 and Ame1-Okp1 and <1% with outer kinetochore subunits (**Figure 5**). Thus, additional factors limit reconstitution relative to the kinetochore *in vivo*, where 1 Cse4 nucleosome recruits 7-8 outer kinetochore complexes [26]. The ability to quantitatively monitor assembly will facilitate developing conditions that improve the reconstitution to match the physiological assembly state.

### Effect of cell cycle on *ex vivo* reconstituted kinetochores

Finally, we addressed the effect of cell cycle stages on kinetochore reconstitution by comparing extracts from G1-phase cells to those from M-phase cells (**Figure 6**). In G1-phase cell extracts, histones associated with DNA (CEN and MUT) robustly (**Figure 6A**). Cbf1, CBF3, as well as other inner kinetochore components reconstituted specifically on CEN-DNA and not MUT-DNA, as observed when using M-phase cell extracts (**S8 Figure**). However, while the assembly of Cbf1 and CBF3 did not show an appreciable difference from M-phase (**Figure 6A**), Cse4, Mif2, and the Ame1-Okp1 complex assembled on CEN-DNA at significantly lower levels in G1-phase cell extracts (**Figure 6B**). Consequently, other kinetochore subunits were reconstituted below the lower limit of detection (**Figure 6C**). These findings suggest that robust assembly of Cse4, besides being the limiting step, is controlled by cell cycle states.

**Figure 6.**
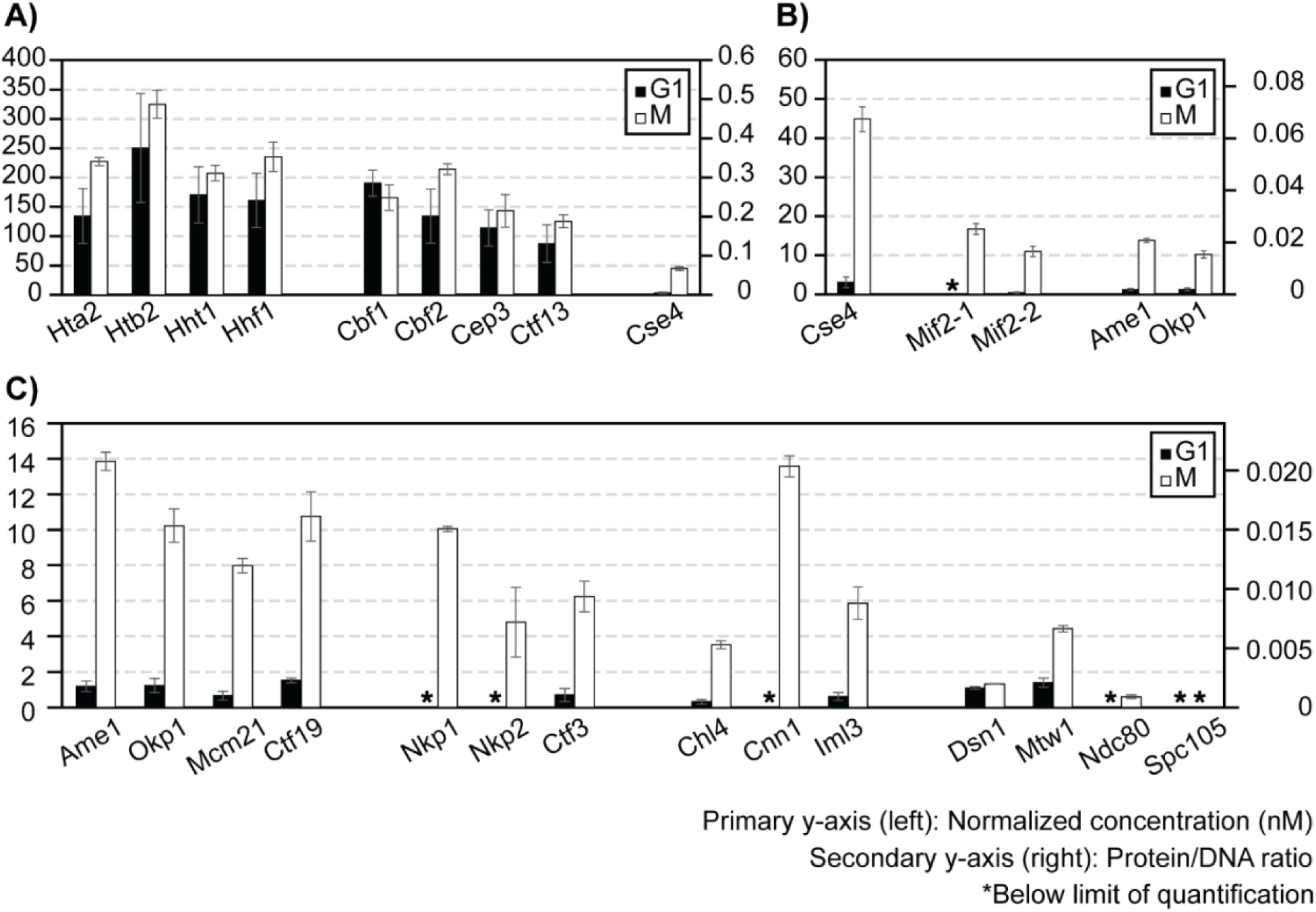
Effect of cell cycle on *ex vivo* reconstituted kinetochores. **A-C)** Quantifications of *ex vivo* reconstituted kinetochore subunits when using G1 or M-phase lysates: y-axes are defined as in Figure 5. The plot shows only the results for CEN-DNA reconstituted kinetochores. See **S8 Figure** for MUT-DNA data of G1-phase reconstituted kinetochores.

## Discussion

In this study, we describe the development and application of Concatemer Assisted Stoichiometry Analysis (CASA), combining LC-PRM-MS and quantitative concatemers derived from a yeast expression system to measure protein complex stoichiometry, determining the reconstitution efficiencies of kinetochore subunits in concentrated yeast cell lysates. The following discussion details the advantages, caveats, and potential improvements of CASA as revealed through this proof-of-concept biological application.

### Design strategies of the CKP

QconCAT and its variants have been used previously to produce quantitative concatemer standards [55,56,70]. Since concatemers are usually unstructured, the expression of CKP in bacteria suffers from protein degradation and the formation of inclusion bodies (unpublished observations) [71]. Therefore, a budding yeast protein expression system was selected and implemented with several optimizations to produce CKP. First, a high-copy expression plasmid with a galactose inducible promoter was used to improve protein yield. Second, an N-terminal GST tag was incorporated to improve CKP solubility and enable absolute quantification (**S3 Figure**). Third, a C-terminal 6xHis-3xFlag tag was used for detection and purification. Fourth, yeast strains available for SILAC were used to introduce heavy isotope-containing amino acids and generate stable isotope-labeled internal standard [57,65].

Selecting peptides with the appropriate length and chemical properties is crucial to minimize challenges during method development and eventual quantification (e.g., poor chromatography, peptide stability, poor ionization, interferences, etc.). Excessively hydrophilic peptides suffer from poor retention during reverse-phase chromatography, while highly hydrophobic peptides are more prone to solubility issues and adsorption losses. For instance, the Htb2 peptide (HAVSEGTR) lacks hydrophobic residues. It binds poorly to the C18 column, leading to suboptimal chromatographic separation (**S7 Figure**). This signal loss, compounded by matrix suppression and interferences, hampered precision when this peptide was analyzed in the reconstituted sample (**Figures 5 and 6**). These issues can be alleviated by choosing a different peptide for quantification or utilizing more internal standards. Hydrophobic peptides, like the Cnn1 peptide (SFLQDLSQVLAR), suffer from stability issues that are likely due to non-specific losses (precipitation/adsorption losses), even in the presence of the stabilization matrix (**Figure 3B and Table 3**). Peptide length is another important factor to consider, as longer peptides provide a greater diversity of transitions to distinguish the target peptide from background noise and interfering ions. Despite these advantages, practical issues should be considered when selecting longer peptides for concatemer proteins. For example, longer peptides would increase the overall length of a concatemer, necessitating the use of multiple concatemers with fewer tryptic peptides in each concatemer to maintain expression and solubility. While using multiple concatemers increases the complexity of analysis, it is not a limiting factor for CASA due to the relative ease of producing full-length concatemers in the yeast expression system.

Finally, our results demonstrate that simultaneous monitoring of multiple peptides for each protein of interest is necessary to ensure quantitative accuracy. In this proof-of-concept study, one peptide was chosen for each kinetochore subunit except for Mif2, for which two peptides were chosen. The observed average difference in the quantitative determinations of the two Mif2 peptides was ∼34% (**Figure 5**). Though modest, this discrepancy exceeds the analytical precision of the method. Thus, a couple of additional factors could have contributed to the discordant quantification: First, flanking amino acids surrounding the tryptic sites of each Mif2 peptide were not included in the CKP, which could affect the trypsin digestion efficiency of CKP versus the native Mif2 protein [72]. Second, as described above, one of the Mif2 peptides (VAPLQYWR) demonstrated greater instability than its counterpart. Such loss could be non-specific and variable for different peptides. There are two approaches to circumvent this possible loss during sample preparation: The first approach is to characterize the loss of each peptide using the CKP; The second, and more robust approach, is to spike the CKP as proteins into the samples before trypsin digestion. In summary, future designs and applications of CKPs should address these limitations, and quantification using more than one peptide specific to each protein should be considered.

### Matrix and CE optimization contribute to optimal analytical performance

The stabilization matrix significantly reduced the recovery losses observed in the more hydrophobic peptides (**Figure 3B** and **Table 3**). As peptide recovery losses were non-linear with respect to time, peptide precipitation or non-specific adsorption loss in-vial was the most likely cause of poor recovery (**Figure 3**). Specifically, matrix stabilization effectively eliminated peptide losses for all except the Cnn1 peptide. As expected, isotope dilution appropriately compensates for the recovery losses of Cnn1, as quantification of the Cnn1 peptide demonstrated acceptable accuracy and precision when isotope diluted, highlighting the need for the utilization of stable isotope-labeled internal standard to ensure robust quantification. Interferences from the stabilization matrix were also evaluated, and little to no interference was observed for PRM scans monitoring the light or the heavy peptides of the CKP. The Htb2, Hht1, Hhf1, and Cbf1 peptides have minimal matrix contribution in the light channel with peak areas less than or comparable (Htb2) to the lowest calibrator. Therefore, incorporating a stabilization matrix aided in the overall robustness of the method and did not interfere with the quantification of the 25 targeted peptides. It’s essential to include the appropriate matrix in CASA as a standard practice to ensure consistent peptide recovery over time.

CE optimization improved the method’s sensitivity for most peptides in the CKP (**Figure 2C**). However, when applied to the analysis of reconstituted kinetochores, interferences were observed for the most abundant product ions for almost all peptides. This stems from specific matrix effects related to reconstitutions performed with CEN, MUT, and ARS DNA; each of these reconstitutions has the potential to have interfering ions unique to the sample. Such occurrences are unavoidable in biological applications and were addressed by changing quantifier ions to those free of interferences.

Characterizing optimal CEs for all product ions of each peptide through PRM allowed us to choose optimal CEs in response to changes in quantifier ions. Use of ion ratios adds further robustness by systematically identifying unknown interferences. Though selecting a quantifier with a lower signal-to-noise ratio impacts analytical sensitivity, specificity is paramount to ensure the accuracy of the quantitative determination.

### Lessons learned about the kinetochore and future directions

Reconstituted kinetochores using centromeric DNA and concentrated cell extracts are the best-known examples of native centromeric DNA successfully wrapped with Cse4, Mif2, CBF3, and all other essential kinetochore subunits [42]. However, due to the relatively low abundance of kinetochores in this reconstitution system and the high complexity of the sample derived from cell lysates, the precise stoichiometry of the *ex vivo* reconstituted kinetochore was not determined [42]. This study applies CASA to determine the efficiency of ex vivo reconstituted kinetochores and examine how it is influenced by the cell cycle. Based on our observations, canonical histones and DNA proximal factors associated with CEN DNA robustly. The equal loading of canonical histone H3 (Hht1) with the other canonical histones (Hta2, Htb2, Hhf1) indicates that the Cse4 loading observed was essentially within the error of canonical histone loading. The inefficient loading of Cse4 in cell lysates was highlighted by the observation that Cse4 : Cbf2 or Cse4 : Cep3 ratios were less than 1 : 1, suggesting that Cse4 assembly may be a limiting step in *ex vivo* kinetochore assembly (Figures 5 and 6). To understand why Cse4 is limiting, follow-up studies evaluating the role of Scm3, the chaperone for Cse4, should be performed [38–40]. Moreover, the cell cycle dependence of Cse4-loading activity was evident, as approximately 14-fold less Cse4 was recovered on centromeric DNA using G1-phase cell extracts relative to M-phase cell extracts, suggesting the involvement of unknown M phase signals in promoting Cse4 loading, in line with this step being limiting in this system (Figure 6).

Overall, the inner kinetochore subunits assembled on centromeric DNA more efficiently than the outer kinetochore subunits (**Figure 5**). This progressive reduction in reconstitution efficiency observed from nucleosome to the outer kinetochore shows that the *ex vivo* reconstituted kinetochore could be missing factors important for stabilizing the complete kinetochore at each stage of its assembly. Future studies utilizing the kinetochore reconstitution system can be complemented with purified proteins to evaluate missing/limiting factors. [45] Such complementation experiments and their evaluation by CASA are essential for understanding the principles of kinetochore assembly.

In summary, this study demonstrates the proof-of-concept application of CASA in studying the efficiency of *ex vivo* reconstituted kinetochores, revealing its potential as a quantitative platform to study native protein complexes in a variety of biological processes.

## Acknowledgment

We thank all members of the Zhou and Suhandynata laboratories at the University of California, San Diego. We thank Dr. Arshad Desai for critical reading of this manuscript, Dr. Swathi Krishnan at Pfizer and the Cell Signaling San Diego group for mentoring JC.

This work was supported by National Institutes of Health GM151191 to R. Suhandynata, GM151191, GM116897 and OD023498 to H. Zhou. JC is additionally supported by the Pfizer-Cell Signaling San Diego Fellowship.

---

**S1 Figure.**
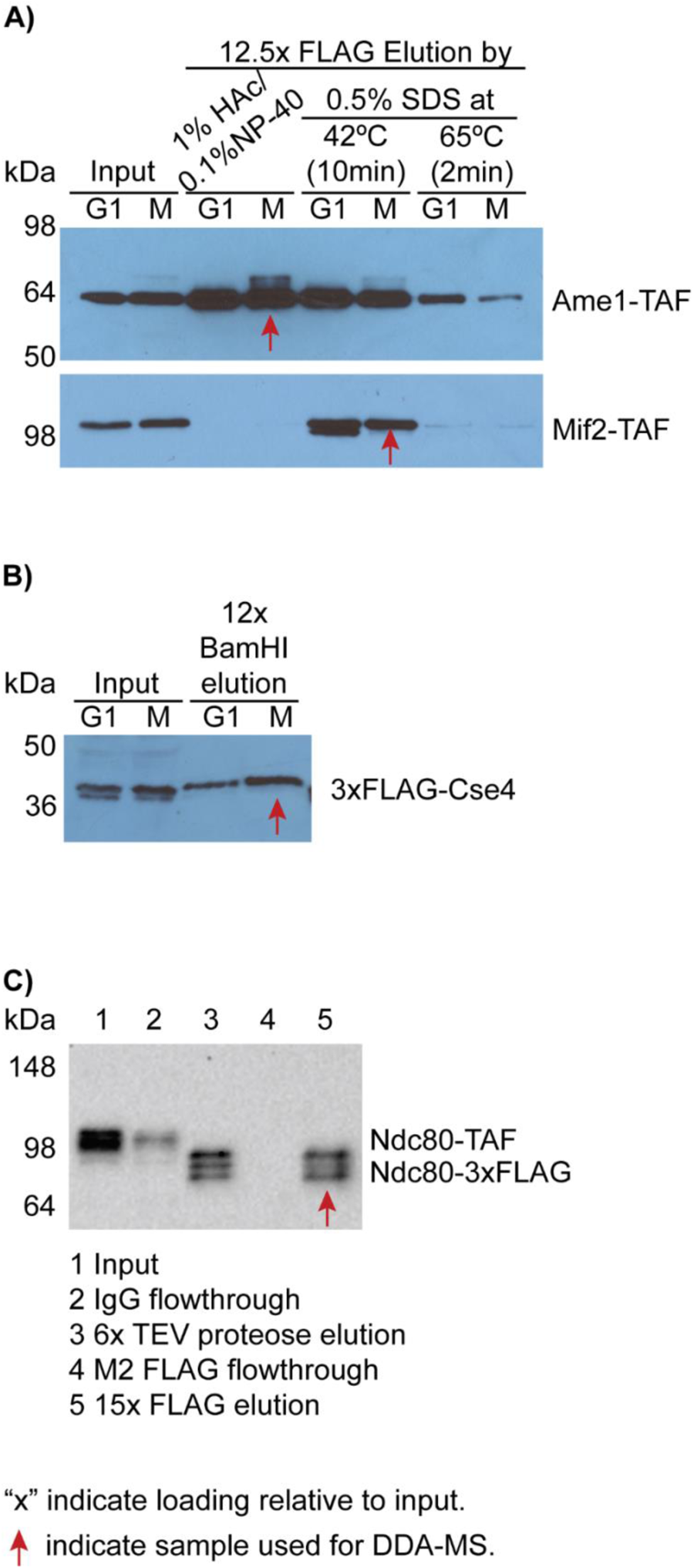
Western blot analysis the immuno-precipitation of TAF-tagged kinetochore subunits. TAF tag: 3xFlag-tev-ProteinA. The purified samples were then processed and analyzed by data-dependent acquisition MS analysis. Red arrows indicate the samples processed for DDA-MS. **A)** M2 FLAG purification of Ame1-TAF and Mif2-TAF from α factor (G1) or Cdc20 auxin degradation (M) arrested cells. HAc stands for acetic acid. (Anti-protein A Western blots) **B)** *Ex vivo* reconstitution of G1 or M phase arrested 3xFLAG-Cse4 strain. (Anti-FLAG Western blot) **C)** 2-step purification of Ndc80 from M phase arrested Ndc80-TAF strain. The first step was done using IgG beads, followed by TEV protease cleavage to cut at the TEV site between protein A and 3xFLAG. The second step was done using M2 FLAG beads, followed by elution with 1% acetic acid/0.1% NP-40. (Anti-FLAG Western blot)

**S2 Figure.**
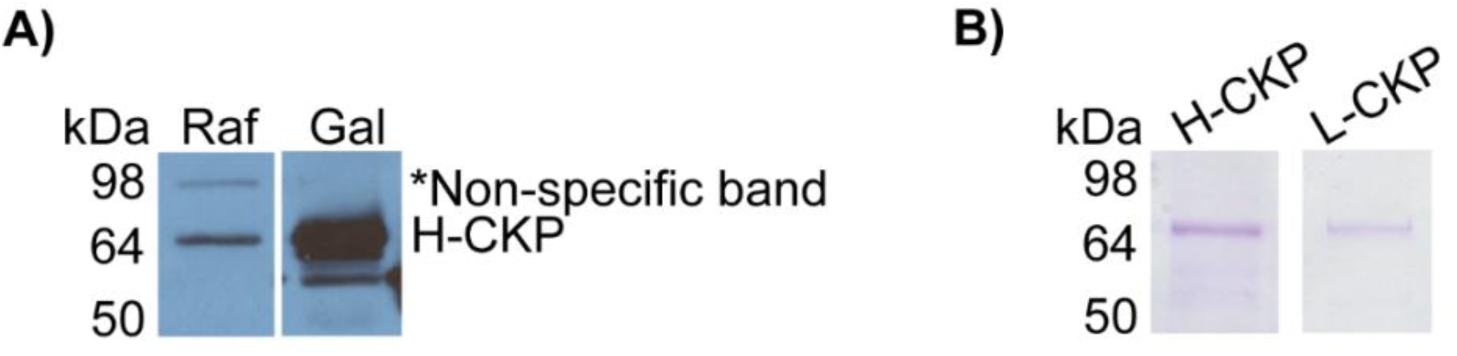
Evaluate CKP expression in yeast by anti-Flag WB and purification via anti-Flag resins. **A)** Anti-FLAG Western blot for galactose induction of H-CKP. Raf: raffinose, Gal: galactose. **B)** Coomassie brilliant blue staining of H and L-CKP after purification. Staining for L and H-CKP was done on different days on different gels.

**S3 Figure.**
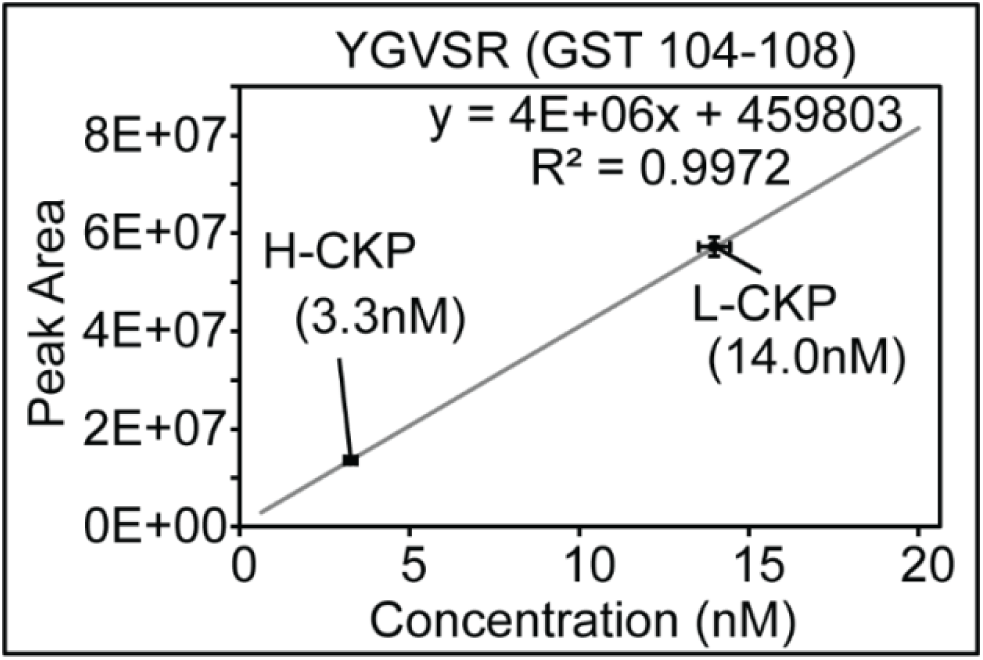
Absolute quantification of L and H-CKP using a commercially synthesized GST peptide (sequence YGVSR) via external calibration.

**S4 Figure.**
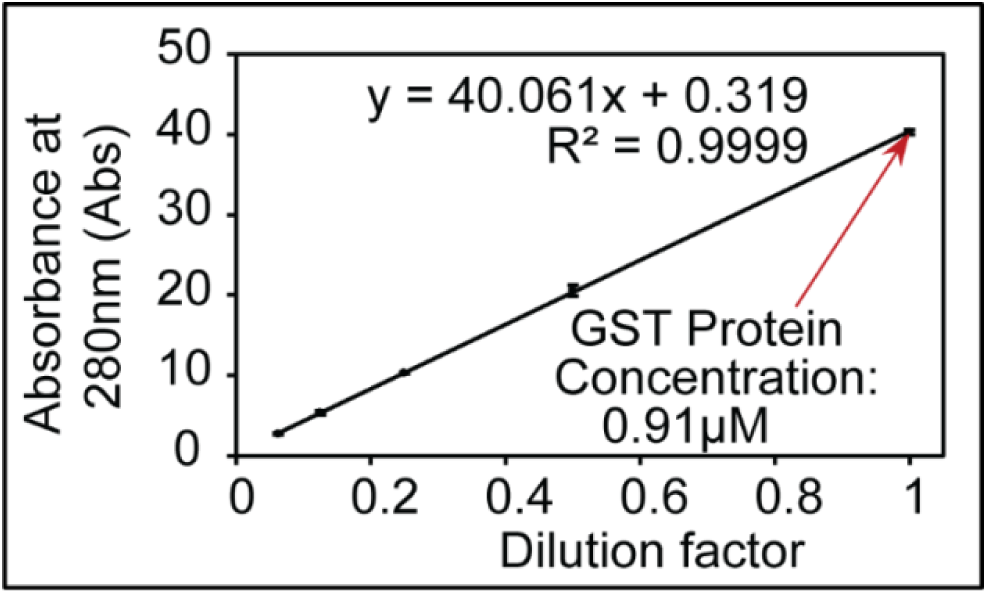
The concentration of recombinant GST protein determined by UV-Vis absorption, which was then used to quantify H-CKP and L-CKP by MS.

**S5 Figure.**
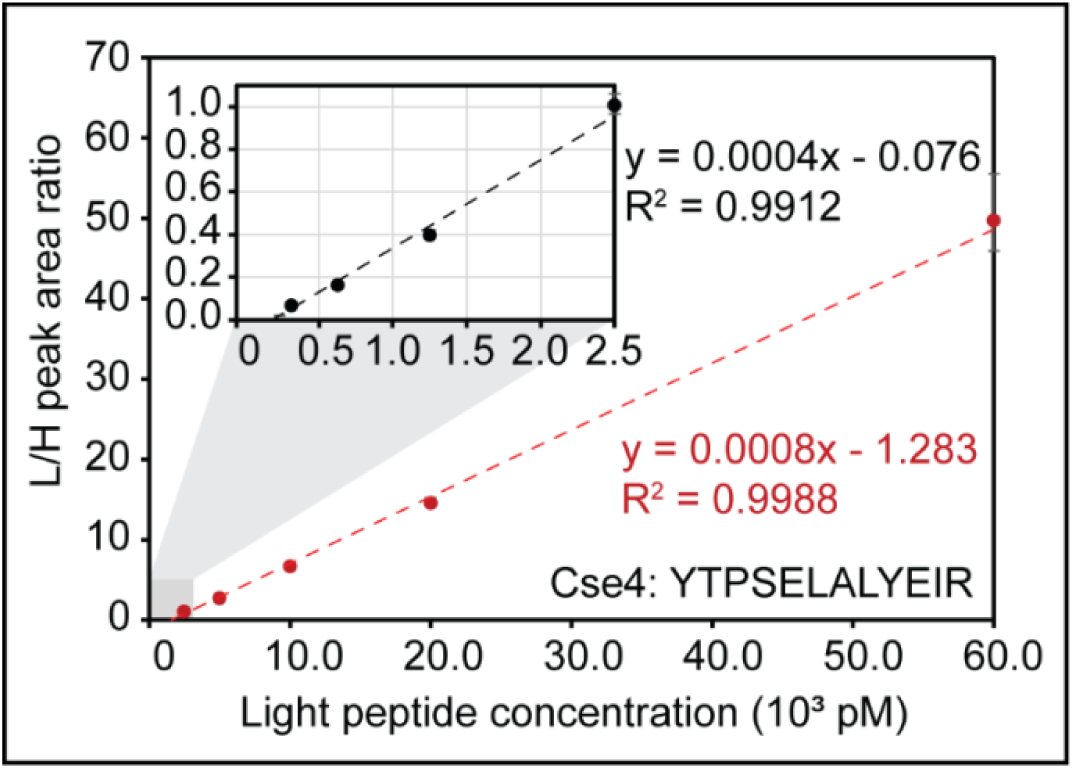
Cse4 has 2 calibration curves covering its full AMR, one at a lower concentration below 2.5 nM, one at the higher concentration up to 60 nM.

**S6 Figure.**
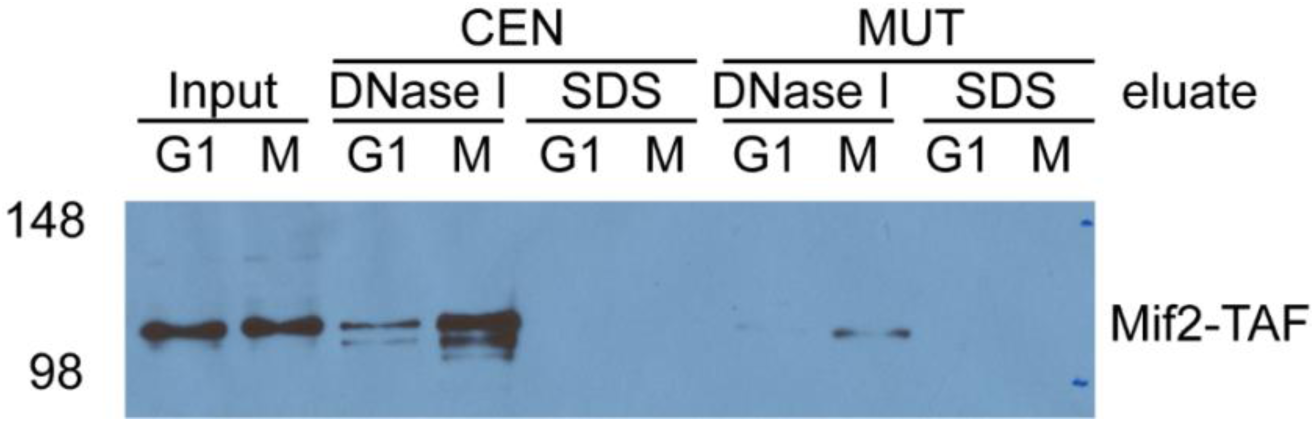
Evaluation of *ex vivo* kinetochore reconstitution using Mif2-TAF cell extracts. M-phase extract has a stronger Mif2-CEN association than G1 extract. Moreover, DNase-I elution is complete in eluting Mif2 since the subsequent SDS elution yielded much fewer signals. Mif2-CEN association is also specific and reduced by point mutation to the CDE III region in CEN3 (MUT).

**S7 Figure.**
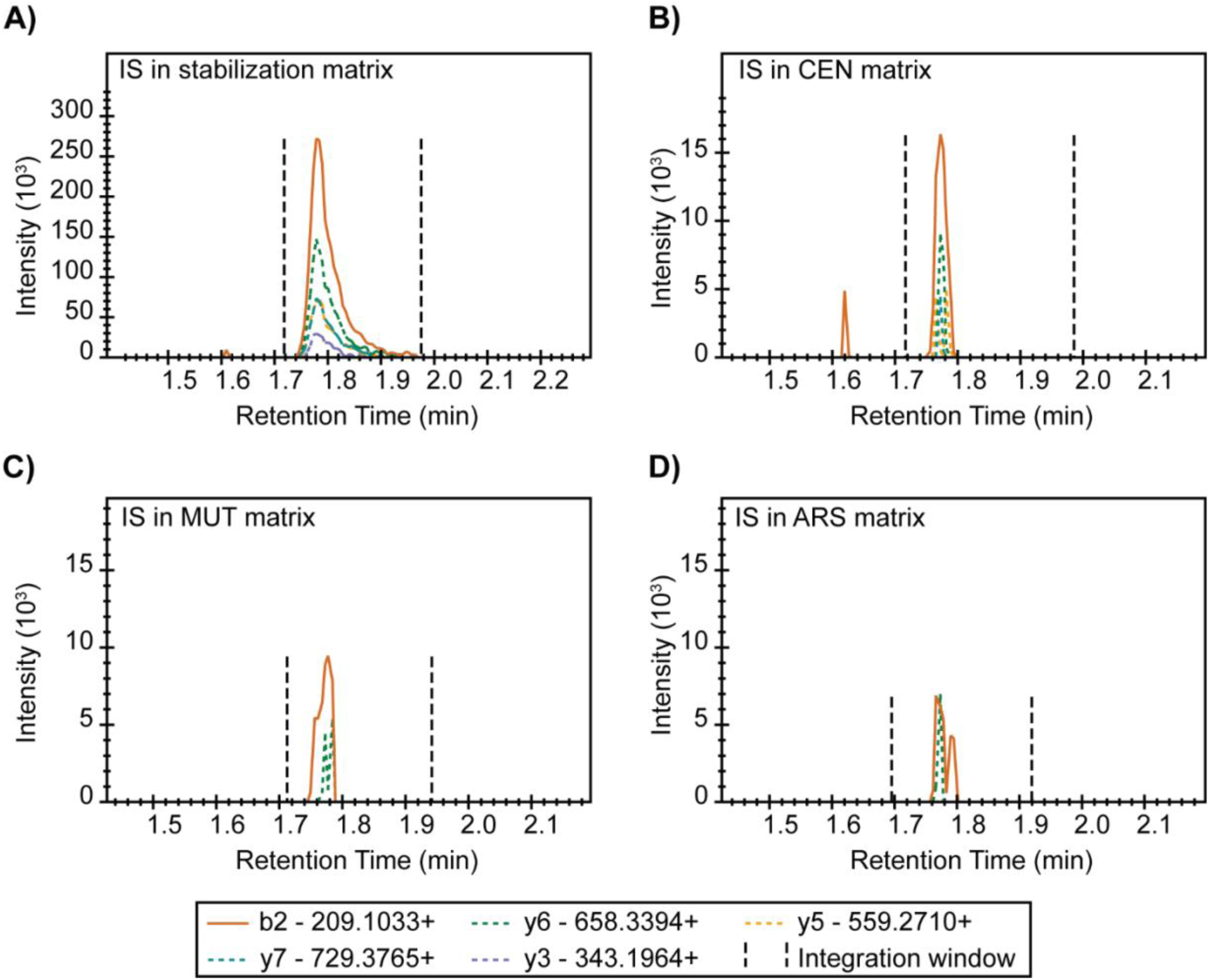
Htb2 peptide chromatograph shows sample-specific matrix suppression. IS: Internal standard, referring to Htb2 peptide here. Chromatograms of indicated transition ions of the Htb2 peptide (HAVSEGTR) from the H-CKP. Y-axes indicate signal intensity, and x-axes indicate retention time in minutes. Black dotted lines indicate the integration windows for determining the peak area of each transition ion.

**S8 Figure.**
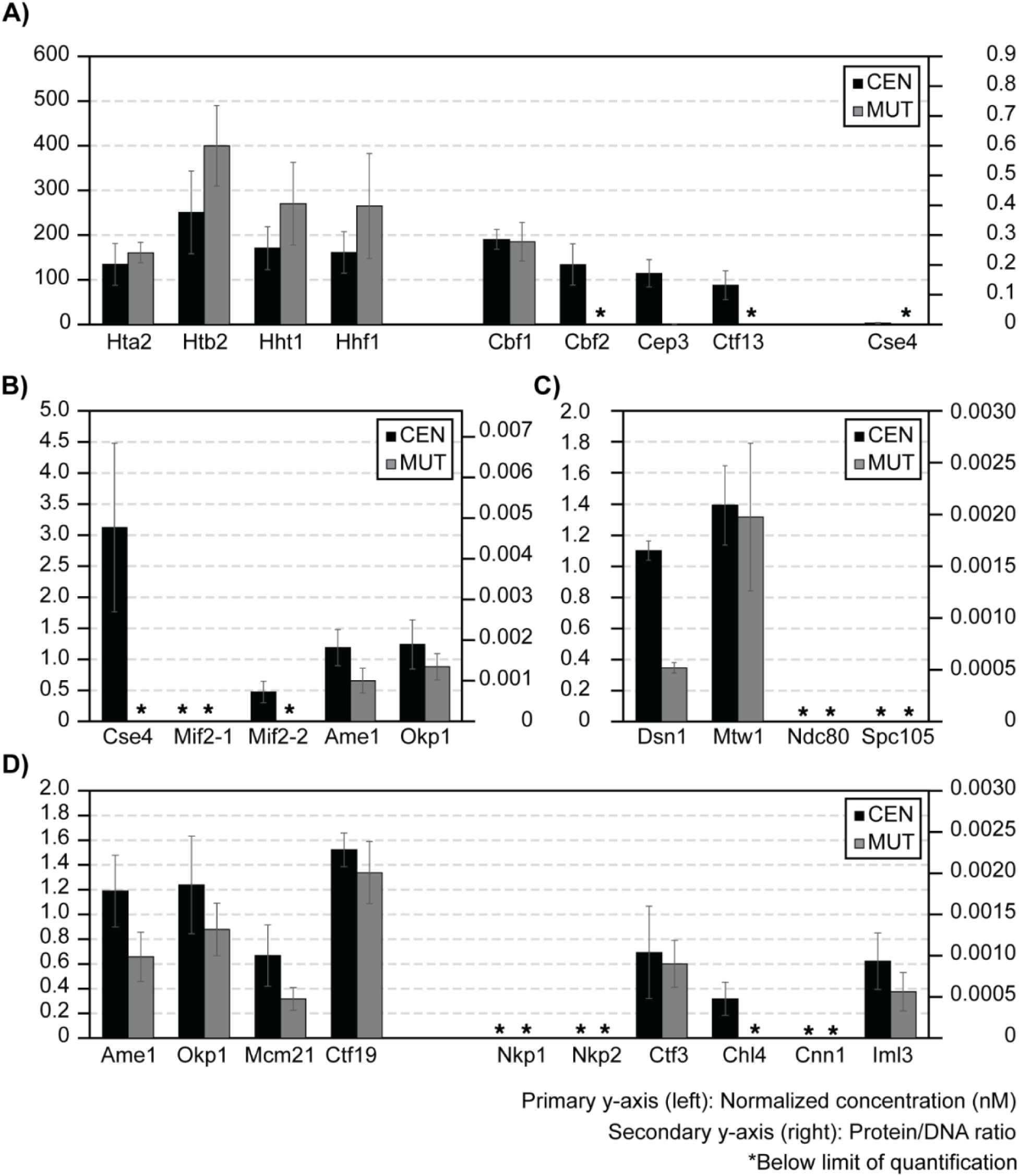
*Ex vivo* reconstitution of kinetochores using G1-phase lysate.

**S1 Table.** Plasmids and yeast strains.

| **Plasmid** | **Description/source** |
| --- | --- |
| HZE2029 | Lic-2GT from Corbett Lab |
| HZE3236 | pRS423-pGAL-GST-tev-HF |
| HZE3240 | Addgene 30116, pFastBacCEN3e2_Plus60_tetO2, from Hinshaw: pSMH1717 |
| HZE3241 | Addgene 30116, pFastBac LIC cloning vector, CEN3e2_Plus60-TetO2-CCGmut, from Hinshaw: pSMH1726 |
| HZE3246 | CEN3-8xLacO-TRP1, from Biggins: pSB964 |
| HZE3361 | pRS423-pGAL-GST-CKP-HF |

| **Yeast Strain** | **Genotype** |
| --- | --- |
| SCY249 | *ura3-52 leu2-1 trp1-63 his3-200 lys2-Bgl hom3-10 ade2-1 ade8 sml1Δ::TRP arg4Δ MAT a* |
| HZY1029 | *bar1Δ::URA MAT a*, W303 in HZY1077 |
| HZY1077 | *ade2-1 can1-100 his3-11,15 leu2-3,112 trp1-1 ura3-1 RAD5+*, MAT a, W303 |
| HZY2347 | *Cdc20-HA-AID::G418 pGPD1-OsTir1::LEU2 bar1Δ::URA Mif2-TAF::HisMX sml1Δ::TRP*, MAT a in SCY249 |
| HZY2461 | *Cdc20-HA-AID::G418 pGPD1-OsTir1::LEU2 bar1Δ::URA Ndc80-TAF::G418 sml1Δ::TRP*, MAT a in SCY249 |
| HZY2646 | *Ame1-TAF::HisMX Cdc20-HA-AID::G418 pGPD1-OsTir1::LEU2 bar1Δ::URA sml1Δ::TRP*, MAT a in SCY249 |
| HZY2777 | *G418::3xFlag-Cse4 Cdc20-HA-AID::HisMX pGPD1-OsTir1::LEU2 bar1Δ::URA*, *sml1Δ::TRP*, MAT a in SCY249 |
| HZY3059 | HZE3361 in SCY249 |

**S2 Table.** Peptides identified by Data-Dependent Acquisition MS (See Supplementary Excel file)

**S3 Table.** Quantification of light and heavy CKP concentrations using a GST peptide standard.

| Samples Analyzed | Synthetic GST peptide: YGVSR |  |  |  |  |  | Light CKP |  |  | Heavy CKP |  |  |
| --- | --- | --- | --- | --- | --- | --- | --- | --- | --- | --- | --- | --- |
| Cal. Levels/<br>Repeat | 1 | 2 | 3 | 4 | 5 | 6 | 1 | 2 | 3 | 1 | 2 | 3 |
| Conc. (pM) | 625 | 1250 | 2500 | 5000 | 10000 | 20000 | 14491 | 13881 | 13564 | 3073.3 | 3302.9 | 3427.6 |
| Peak area (10<sup>3</sup>) | 2567 | 5746 | 10155 | 23086 | 38492 | 82090 | 59331 | 56842 | 55550 | 12804 | 13740 | 14247 |
| Avg. conc. (pM) |  |  |  |  |  |  | 13978.67 |  |  | 3267.93 |  |  |
| Avg. peak area (10<sup>3</sup>) |  |  |  |  |  |  | 57241 |  |  | 13597 |  |  |
| Std. dev conc. |  |  |  |  |  |  | 471.15 |  |  | 179.72 |  |  |
| Std. dev peak area (10<sup>3</sup>) |  |  |  |  |  |  | 1921 |  |  | 732 |  |  |

**S4 Table.** Target inclusion list for PRM-MS (See supplementary Excel file)

**S5 Table.** Isotope incorporation efficiencies of H-CKP. The peak areas shown are the sum of the peak areas of the top 12 transitions.

| **Peptide** | **Heavy area** | **Light area** | **% Heavy of total area** |
| --- | --- | --- | --- |
| HTA2 | 17078343 | 0 | 100.00 |
| CTF3 | 20109657 | 0 | 100.00 |
| CBF2 | 19143204 | 1570 | 99.99 |
| NKP1 | 22143101 | 20836 | 99.91 |
| IML3 | 20163754 | 0 | 100.00 |
| HTB2 | 8778000 | 10597 | 99.88 |
| MCM21 | 32181955 | 1491 | 100.00 |
| DSN1 | 38273619 | 0 | 100.00 |
| MTW1 | 40463111 | 0 | 100.00 |
| HHF1 | 24330620 | 0 | 100.00 |
| CBF1 | 67708609 | 57839 | 99.91 |
| CEP3 | 22579733 | 0 | 100.00 |
| AME1 | 14451112 | 0 | 100.00 |
| CHL4 | 44155600 | 62964 | 99.86 |
| CTF19 | 27480906 | 1417 | 99.99 |
| NDC80 | 26923566 | 0 | 100.00 |
| CNN1 | 13304552 | 1245 | 99.99 |
| HHT1 | 29087574 | 0 | 100.00 |
| CTF13 | 28351430 | 0 | 100.00 |
| MIF2-2 | 30827943 | 105111 | 99.66 |
| SPC105 | 16052332 | 0 | 100.00 |
| OKP1 | 10856738 | 0 | 100.00 |
| NKP2* | 18603646 | 391167* | 97.94* |
| MIF2-1 | 41905182 | 0 | 100.00 |
| CSE4 | 56761875 | 2634 | 100.00 |

\* Most abundant transition (b2) is interfered with by a contaminant.
Actual isotope incorporation > 99%

**S6 Table.** Accuracy of calibrators (Tolerance: ± 20%): Gray cells are outside the AMR (Tolerance: ± 20%). Cells with yellow highlights have %biases between -20% and -15% or 15% and 20%. The rest of the cells have %biases within -15% and 15%.

| Peptide | Calibrator %Bias |  |  |  |  |  |  |  |  |  |
| --- | --- | --- | --- | --- | --- | --- | --- | --- | --- | --- |
| Ame1 | 18.0 | 5.2 | -1.2 | 1.9 | -10.0 | -4.6 | -3.5 | -3.9 | -5.0 | 3.0 |
| Cbf1 | 7.7 | 9.2 | -1.6 | -0.7 | -2.5 | -0.6 |  | -6.9 | -8.7 | 4.1 |
| Cbf2 |  |  |  | -3.3 | -6.0 | 7.2 | 8.9 | 3.3 | -13.0 | 2.9 |
| Cep3 | 3.3 | 3.1 | -5.9 | 8.4 | -1.2 | -2.0 | 1.7 | 3.3 | -15.1 | 4.4 |
| Chl4 | -16.3 | -5.8 | -4.7 | 3.3 | 5.2 | 10.5 | 11.5 | 0.6 | -4.1 | -0.2 |
| Cnn1 |  |  |  |  |  | 15.9 | -14.4 | -6.2 | 4.7 |  |
| Cse4 |  |  | 11.0 | -7.3 | -8.4 | 4.6 |  |  |  |  |
| Ctf13 |  | 0.8 | -8.0 | 2.1 | 0.2 | 1.0 | 5.8 | -0.7 | -1.2 |  |
| Ctf19 |  | 6.5 | 2.6 | 0.5 | 2.3 | -12.5 | 3.9 | -4.1 |  | 0.8 |
| Ctf3 | 2.2 | -1.1 | -8.6 | 4.1 | -1.8 | -0.1 | 6.5 | -0.9 |  | -0.4 |
| Dsn1 | 8.7 | -1.9 | -2.6 | -2.5 | -7.3 | -1.5 | 6.2 | 1.5 |  | -0.5 |
| Hhf1 |  |  |  |  | 13.4 | -7.4 | -0.5 | -6.7 |  | 1.2 |
| Hht1 |  |  | 0.8 | 3.2 | 2.7 | -2.6 | -3.1 | -1.0 | -0.6 | 0.6 |
| Hta2 | 0.7 | 0.7 | -2.9 | 8.4 | -0.1 | -4.0 | 1.5 | -5.1 |  | 0.8 |
| Htb2 |  |  |  | 2.8 | -1.6 | 3.1 | -1.5 | -2.3 | -1.4 | 0.8 |
| Iml3 | 0.4 | 0.1 | -4.9 | 11.0 | -3.5 | 2.6 | -1.4 | -0.3 | -6.1 | 2.1 |
| Mcm21 | -3.9 | -3.5 | 3.4 | 5.8 | -1.7 | -1.1 | 5.5 | -2.0 | -3.5 | 1.0 |
| Mif2-1 |  |  |  | -9.1 | -1.1 | 4.2 | 1.8 | 5.3 |  | -1.1 |
| Mif2-2 | 14.3 | -1.5 | 7.6 | 4.8 | -4.3 | -14.1 |  | -8.7 |  | 2.0 |
| Mtw1 |  |  | 13.7 | -13.1 | -4.3 | 3.7 |  |  |  |  |
| Ndc80 | 1.9 | 19.2 | -9.8 | -1.0 | -1.9 | -5.3 | 0.1 | -4.0 |  | 0.9 |
| Nkp1 |  |  |  | -1.8 | -6.1 | 9.1 | 5.5 | 2.8 | -12.6 | 3.0 |
| Nkp2 |  |  | -8.8 | -3.9 | 1.0 | 6.5 | 4.0 | 3.7 | -2.2 | -0.4 |
| Okp1 |  | -11.5 | 2.6 | 8.5 | 0.2 | 4.7 | 0.4 | -0.8 | -5.6 | 1.7 |
| Spc105 |  |  | 1.6 | -4.8 | -1.3 | -0.5 | 3.6 | 2.0 |  | -0.5 |
| Cse4 - High range |  |  |  |  |  | 10.1 | -3.4 | -4.0 | -4.9 | 2.2 |
| Mtw1 - High range |  |  |  |  |  | 0.1 | 0.1 | -0.3 | 0.1 |  |
| Analyte conc. (pM) | 78 | 156 | 313 | 625 | 1250 | 2500 | 5000 | 10000 | 20000 | 60000 |

**S7 Table.** Inter-day accuracy of calibrators (Tolerance: ± 20%): Gray cells are outside the AMR (Tolerance: ± 20%). Cells with yellow highlights have average %biases between -20% and -15% or 15% and 20%. The rest of the cells have average %biases within -15% and 15%.

| Peptide | Calibrator Avg. %Bias (n = 3) |  |  |  |  |  |  |  |  |  |
| --- | --- | --- | --- | --- | --- | --- | --- | --- | --- | --- |
| Ame1 | 19.5 | 6.8 | 0.3 | 1.8 | -9.9 | -1.6 | -1.7 | -1.6 | -3.5 | 3.5 |
| Cbf1 | 10.2 | 7.7 | -0.7 | 3.1 | -3.0 | -5.9 |  | 0.5 | -4.1 | 2.0 |
| Cbf2 |  |  |  | -7.3 | -5.4 | 3.3 | 8.2 | 4.8 | -7.6 | -0.5 |
| Cep3 | 2.4 | 8.5 | 2.9 | 7.1 | 1.4 | -1.4 | -2.5 | 1.7 | -12.0 | 6.6 |
| Chl4 | -12.7 | -2.1 | 1.6 | 10.5 | 7.1 | 12.8 | 9.4 | 3.5 | -5.8 | 1.4 |
| Cnn1 |  |  |  |  |  | 16.6 | -14.2 | -4.3 | 12.6 |  |
| Cse4 |  |  | 11.9 | -12.1 | -5.9 | 5.3 |  |  |  |  |
| Ctf13 |  | -4.1 | 5.7 | 8.8 | -0.5 | 6.9 | 1.9 | 0.8 | -2.7 |  |
| Ctf19 |  | 9.9 | 4.3 | -6.9 | -1.5 | -11.9 | 1.0 | 0.3 |  | 2.3 |
| Ctf3 | 9.1 | 2.6 | -0.2 | -1.0 | -0.8 | -1.4 | 4.2 | 0.7 |  | -2.0 |
| Dsn1 | 2.7 | -6.8 | -2.6 | -7.8 | -3.1 | -11.4 | 7.6 | 0.0 |  | 4.0 |
| Hhf1 |  |  |  |  | 15.8 | -7.8 | -3.3 | -2.3 |  | 1.5 |
| Hht1 |  |  | -9.4 | 10.5 | -0.1 | -2.3 | -1.7 | -3.9 | -3.8 | 3.7 |
| Hta2 | 2.3 | 6.5 | 0.4 | 4.6 | 0.3 | -1.0 | -1.3 | -2.1 |  | -0.7 |
| Htb2 |  |  |  | 7.4 | -3.2 | 1.8 | 2.2 | 2.0 | -2.1 | -1.2 |
| Iml3 | -10.2 | -1.7 | -0.3 | 4.5 | -3.6 | -0.4 | -2.4 | -8.3 | -7.5 | -0.3 |
| Mcm21 | -5.6 | -2.2 | 1.7 | 5.1 | -5.7 | 1.0 | 0.0 | -5.7 | -5.4 | 6.6 |
| Mif2-1 |  |  |  | -16.2 | -5.3 | -3.6 | 5.3 | 5.3 |  | 3.0 |
| Mif2-2 | 7.9 | -2.8 | 7.8 | -7.0 | -0.5 | -13.7 |  | -5.4 |  | -5.2 |
| Mtw1 |  |  | 15.1 | -15.3 | -10.4 | -0.8 |  |  |  |  |
| Ndc80 | -15.0 | -6.6 | -3.9 | -5.2 | -0.8 | -9.1 | 6.1 | -5.9 |  | -0.4 |
| Nkp1 |  |  |  | -11.0 | 1.8 | 1.2 | 6.4 | 1.4 | -10.3 | 5.0 |
| Nkp2 |  |  | -4.1 | 10.2 | -0.8 | 3.2 | 5.5 | 4.7 | 5.2 | 11.0 |
| Okp1 |  | -18.0 | 0.5 | 11.3 | 1.2 | 3.8 | 2.5 | -3.1 | 0.7 | 2.4 |
| Spc105 |  |  | -0.3 | 1.0 | -4.1 | -3.1 | 7.2 | -3.1 |  | 4.8 |
| Cse4 - High range |  |  |  |  |  | 10.5 | -3.3 | -6.0 | -1.8 | -0.6 |
| Mtw1 - High range |  |  |  |  |  | -2.5 | 2.2 | 6.1 | 10.7 |  |
| Analyte conc. (pM) | 78 | 156 | 313 | 625 | 1250 | 2500 | 5000 | 10000 | 20000 | 60000 |

**S8 Table.** Inter-day precision of calibrators (Tolerance: ± 15%): Gray cells are outside the AMR (Tolerance: ± 15%). Cells with yellow highlights have %CVs between -15% and -10% or 10% and 15%. The rest of the cells have %CVs within -10% and 10%.

| Peptide | Calibrator %CV (n = 3) |  |  |  |  |  |  |  |  |  |
| --- | --- | --- | --- | --- | --- | --- | --- | --- | --- | --- |
| Ame1 | 3.6 | 2.9 | 1.7 | 5.8 | 2.6 | 3.2 | 0.7 | 2.4 | 1.0 | 0.6 |
| Cbf1 | 4.6 | 5.5 | 5.0 | 2.2 | 0.7 | 5.0 |  | 4.0 | 2.7 | 6.6 |
| Cbf2 |  |  |  | 6.5 | 5.0 | 4.8 | 5.3 | 1.8 | 2.2 | 1.7 |
| Cep3 | 11.2 | 0.6 | 1.1 | 3.2 | 1.8 | 9.9 | 1.9 | 0.1 | 0.8 | 9.9 |
| Chl4 | 8.0 | 6.5 | 7.9 | 3.9 | 5.1 | 6.0 | 2.0 | 1.7 | 1.7 | 1.1 |
| Cnn1 |  |  |  |  |  | 1.3 | 1.4 | 5.1 | 7.7 |  |
| Cse4 |  |  | 2.6 | 0.9 | 2.4 | 4.3 |  |  |  |  |
| Ctf13 |  | 1.9 | 5.4 | 2.6 | 0.2 | 4.2 | 2.3 | 6.8 | 3.3 |  |
| Ctf19 |  | 3.1 | 3.2 | 2.3 | 2.5 | 1.3 | 4.7 | 2.4 |  | 0.5 |
| Ctf3 | 14.5 | 4.2 | 2.0 | 6.4 | 5.2 | 2.4 | 4.3 | 3.1 |  | 4.3 |
| Dsn1 | 14.1 | 5.0 | 0.8 | 5.2 | 4.7 | 4.1 | 5.9 | 4.2 |  | 2.6 |
| Hhf1 |  |  |  |  | 2.5 | 1.0 | 2.3 | 1.3 |  | 2.6 |
| Hht1 |  |  | 5.3 | 10.6 | 3.7 | 1.7 | 2.1 | 6.2 | 7.2 | 4.5 |
| Hta2 | 3.4 | 6.0 | 3.7 | 6.8 | 4.7 | 1.9 | 2.4 | 1.6 |  | 1.0 |
| Htb2 |  |  |  | 2.3 | 2.6 | 4.3 | 2.6 | 2.9 | 2.9 | 5.3 |
| Iml3 | 8.5 | 4.5 | 1.4 | 4.7 | 9.5 | 2.5 | 2.9 | 4.1 | 3.2 | 0.9 |
| Mcm21 | 10.8 | 2.2 | 1.4 | 0.1 | 0.4 | 4.1 | 2.7 | 1.2 | 0.5 | 2.6 |
| Mif2-1 |  |  |  | 6.8 | 5.3 | 3.6 | 4.2 | 7.6 |  | 3.8 |
| Mif2-2 | 8.6 | 7.5 | 2.1 | 5.9 | 7.4 | 5.5 |  | 5.5 |  | 3.5 |
| Mtw1 |  |  | 1.9 | 2.2 | 1.0 | 1.5 |  |  |  |  |
| Ndc80 | 7.6 | 10.2 | 9.4 | 5.0 | 4.9 | 2.8 | 5.3 | 10.0 |  | 3.4 |
| Nkp1 |  |  |  | 2.6 | 4.0 | 3.2 | 7.3 | 8.1 | 8.4 | 6.5 |
| Nkp2 |  |  | 5.3 | 6.1 | 1.7 | 7.8 | 4.4 | 5.8 | 3.7 | 5.3 |
| Okp1 |  | 5.8 | 5.2 | 0.9 | 1.7 | 0.5 | 2.3 | 5.5 | 2.8 | 2.8 |
| Spc105 |  |  | 6.4 | 6.1 | 12.9 | 3.5 | 0.6 | 2.8 |  | 4.6 |
| Cse4 - High range |  |  |  |  |  | 2.0 | 1.2 | 1.4 | 1.0 | 0.9 |
| Mtw1 - High range |  |  |  |  |  | 0.9 | 1.0 | 4.2 | 5.1 |  |
| Analyte conc. (pM) | 78 | 156 | 313 | 625 | 1250 | 2500 | 5000 | 10000 | 20000 | 60000 |

**S9 Table.** Quantifier and qualifier ions (L-CKP): Quantifier and qualifier transition ions for the L-CKP and corresponding ion ratios as determined from the average of the calibrators.

| Peptide | Transitions |  |  |  | Quant/Qual ion ratio |
| --- | --- | --- | --- | --- | --- |
|  | Quantifier | Quantifier m/z | Qualifier | Qualifier m/z |  |
| Cbf1 | T - y10++ | 593.7704 | S - y11++ | 637.2864 | 1.33 |
| Cep3 | Y - y5+ | 681.3566 | T - y3+ | 405.2092 | 5.62 |
| Ctf13 | D - y4+ | 538.2620 | L - b3+ | 272.1605 | 1.10 |
| Cse4 | A - y6+ | 764.4301 | L - y5+ | 693.3930 | 1.45 |
| Htb2 | A - b2+ | 209.1033 | V - y6+ | 648.3311 | 1.94 |
| Hta2 | P - y4+ | 428.2616 | F - y5+ | 575.3300 | 1.19 |
| Hhf1 | Y - y5+ | 695.3359 | I - y6+ | 808.4199 | 1.19 |
| Hht1 | E - y5+ | 643.4137 | L - y3+ | 401.2871 | 1.18 |
| Mif2-1 | S - y5+ | 545.3042 | S - y7+ | 761.3788 | 1.43 |
| Mif2-2 | L - y5+ | 765.4042 | Q - y4+ | 652.3202 | 0.58 |
| Cbf2 | N - y9+ | 989.4283 | V - y10+ | 1088.4967 | 1.17 |
| Mcm21 | D - b3+ | 344.1452 | D - y7+ | 793.3686 | 2.09 |
| Ctf19 | D - y6+ | 747.3268 | S - y9+ | 1060.5269 | 2.01 |
| Ctf3 | G - y9+ | 958.5680 | I - y4+ | 529.3457 | 0.98 |
| Iml3 | T - y4+ | 464.2463 | V - y5+ | 563.3148 | 1.89 |
| Chl4 | P - y6+ | 734.4196 | D - y10+ | 1122.5426 | 2.74 |
| Mtw1 | P - y11++ | 676.8277 | L - y7+ | 834.4468 | 1.68 |
| Cnn1 | A - y2+ | 246.1561 | L - b10+ | 1131.6045 | 2.64 |
| Nkp1 | E - y3+ | 417.2456 | D - y7+ | 862.3901 | 0.32 |
| Nkp2 | S - y6+ | 704.3573 | L - y4+ | 488.2827 | 1.72 |
| Ndc80 | D - y9+ | 1033.5273 | S - y7+ | 831.4683 | 1.47 |
| Dsn1 | E - y9+ | 1154.4960 | Y - y7+ | 911.4105 | 0.97 |
| Spc105 | Y - y5+ | 609.2991 | S - y4+ | 446.2358 | 0.78 |
| Okp1 | Q - y5+ | 666.3206 | A - y4+ | 538.2620 | 1.20 |
| Ame1 | D - y5+ | 605.3253 | E - y6+ | 734.3679 | 0.81 |

**S10 Table.** Quantifier and qualifier ions (H-CKP)

| Peptide | Transitions |  |  |  | Quant/Qual ion ratio |
| --- | --- | --- | --- | --- | --- |
|  | Quantifier | Quantifier m/z | Qualifier | Qualifier m/z |  |
| Cbf1 | T - y10++ | 598.7745 | S - y11++ | 642.2905 | 1.33 |
| Cep3 | Y - y5+ | 691.3649 | T - y3+ | 415.2175 | 5.62 |
| Ctf13 | D - y4+ | 548.2703 | L - b3+ | 272.1605 | 1.10 |
| Cse4 | A - y6+ | 774.4384 | L - y5+ | 703.4013 | 1.45 |
| Htb2 | A - b2+ | 209.1033 | V - y6+ | 658.3394 | 1.94 |
| Hta2 | P - y4+ | 438.2699 | F - y5+ | 585.3383 | 1.19 |
| Hhf1 | Y - y5+ | 705.3442 | I - y6+ | 818.4282 | 1.19 |
| Hht1 | E - y5+ | 653.4220 | L - y3+ | 411.2953 | 1.18 |
| Mif2-1 | S - y5+ | 555.3125 | S - y7+ | 771.3871 | 1.43 |
| Mif2-2 | L - y5+ | 775.4125 | Q - y4+ | 662.3284 | 0.58 |
| Cbf2 | N - y9+ | 999.4365 | V - y10+ | 1098.505 | 1.17 |
| Mcm21 | D - b3+ | 344.1452 | D - y7+ | 803.3769 | 2.09 |
| Ctf19 | D - y6+ | 757.3350 | S - y9+ | 1070.535 | 2.01 |
| Ctf3 | G - y9+ | 968.5763 | I - y4+ | 539.3539 | 0.98 |
| Iml3 | T - y4+ | 474.2546 | V - y5+ | 573.323 | 1.89 |
| Chl4 | P - y6+ | 744.4278 | D - y10+ | 1132.551 | 2.74 |
| Mtw1 | P - y11++ | 681.8318 | L - y7+ | 844.4551 | 1.68 |
| Cnn1 | A - y2+ | 256.1643 | L - b10+ | 1131.604 | 2.64 |
| Nkp1 | E - y3+ | 427.2539 | D - y7+ | 872.3984 | 0.32 |
| Nkp2 | S - y6+ | 714.3656 | L - y4+ | 498.291 | 1.72 |
| Ndc80 | D - y9+ | 1043.5355 | S - y7+ | 841.4766 | 1.47 |
| Dsn1 | E - y9+ | 1164.5043 | Y - y7+ | 921.4188 | 0.97 |
| Spc105 | Y - y5+ | 619.3074 | S - y4+ | 456.244 | 0.78 |
| Okp1 | Q - y5+ | 676.3288 | A - y4+ | 548.2703 | 1.20 |
| Ame1 | D - y5+ | 615.3336 | E - y6+ | 744.3762 | 0.81 |

**S11 Table.** Precision (%CV) of process replicates (n = 3) of ex vivo kinetochore reconstitutions.

| Peptide | %CV |  |  |  |  |
| --- | --- | --- | --- | --- | --- |
|  | M |  |  | G1 |  |
|  | ARS | CEN | MUT | CEN | MUT |
| **Ame1** | 19.8 | 3.7 | 21.3 | 24.4 | 30.2 |
| **Cbf1** | 78.0 | 13.3 | 10.4 | 11.7 | 23.4 |
| **Cbf2** | 6.2 | 4.4 | 20.4 | 34.4 |  |
| **Cep3** | 15.4 | 19.4 | 10.3 | 26.8 | 50.5 |
| **Chl4** |  | 6.2 |  | 42.2 |  |
| **Cnn1** |  | 4.4 |  |  |  |
| **Cse4** | 5.2 | 7.2 | 6.3 | 43.5 |  |
| **Ctf13** | 12.8 | 8.8 | 30.4 | 37.1 |  |
| **Ctf19** | 9.6 | 12.9 | 12.0 | 8.9 | 18.9 |
| **Ctf3** |  | 13.8 | 20.9 | 53.9 | 31.4 |
| **Dsn1** | 4.0 | 0.7 | 4.7 | 5.6 | 9.5 |
| **Hhf1** | 10.8 | 10.6 | 17.9 | 28.8 | 44.3 |
| **Hht1** | 12.5 | 6.3 | 16.9 | 28.1 | 34.1 |
| **Hta2** | 5.7 | 3.1 | 10.4 | 34.9 | 14.4 |
| **Htb2** | 6.7 | 7.3 | 19.9 | 37.1 | 22.5 |
| **Iml3** |  | 15.5 | 23.6 | 36.6 | 41.2 |
| **Mcm21** |  | 5.0 | 26.9 | 37.2 | 29.0 |
| **Mif2-1** |  | 8.6 |  |  |  |
| **Mif2-2** | 16.5 | 12.5 |  | 36.8 |  |
| **Mtw1** | 10.4 | 4.2 | 11.4 | 18.4 | 36.1 |
| **Ndc80** |  | 17.9 |  |  |  |
| **Nkp1** |  | 1.4 |  |  |  |
| **Nkp2** |  | 40.8 |  |  |  |
| **Okp1** |  | 9.1 | 9.8 | 31.9 | 24.2 |
| **Spc105** |  |  |  |  |  |

%CVs were not calculated for measurements < lower limit of quantification, which are shown as empty cells in the table above.

